# A *Trypanosoma cruzi* Antigen and Epitope Atlas: deep characterization of antibody specificities in Chagas Disease patients across the Americas

**DOI:** 10.1101/2022.08.19.504544

**Authors:** Alejandro D. Ricci, Leonel Bracco, Janine M. Ramsey, Melissa S. Nolan, M. Katie Lynn, Jaime Altcheh, Griselda Ballering, Faustino Torrico, Norival Kesper, Juan C. Villar, Jorge D. Marco, Fernán Agüero

## Abstract

During an infection, the immune system produces pathogen-specific antibodies. With time, these antibody repertoires become specific to the history of infections and represent a rich source of diagnostic markers. However, the specificities of these antibodies are mostly unknown. Here, using high-density peptide arrays we examined the specificities of human antibody repertoires of Chagas disease patients. Chagas disease is a neglected disease caused by Trypanosoma cruzi, a protozoan parasite that evades immune mediated elimination and mounts long-lasting chronic infections. We describe here the first proteome-wide search for antigens and epitopes and their seroprevalence at the individual level and across human populations. In a first discovery screening of 2.84 million short peptides spanning two *T. cruzi* proteomes we found 3,868 distinct antigenic protein regions. Further analysis of repertoires from 71 individuals provided information on their seroprevalence and showed a large fraction of private epitopes of low seroprevalence (<20%), and novel high seroprevalence antigens. Using single-residue mutagenesis we found the core epitopes required for antibody binding for 232 of these epitopes. These datasets enable the study of the Chagas antibody repertoire at an unprecedented depth and granularity, while also providing a rich source of novel serological biomarkers.

**IMPACT STATEMENT:** This work reveals the diversity and extent of antibody specificities in Chagas Disease and provides a wealth of well-defined antigenic markers for diagnosis and development of serological applications for this neglected infectious disease.

## INTRODUCTION

Although the molecular mechanisms that produce diverse antibody repertoires are precisely understood (Di Noia and Neuberger, 2007; Neuberger, 2008), a comprehensive description of their specificities in different infected individuals has been hindered by the lack of powerful tools.

Synthetic peptides have been used historically to map continuous antibody-binding epitopes or to find key residues for protein binding at small scale (Frank and Overwin, 1996; Geysen et al., 1987; Reineke and Sabat, 2009). With the introduction of peptide arrays, it was possible to display large numbers of peptides on a solid surface at addressable positions (Pellois et al., 2002; Vengesai et al., 2022). Given the sustained increase in the densities achieved by the *in situ* synthesis of peptides using maskless photolithography (Buus et al., 2012; Legutki et al., 2014; Carmona et al., 2015; Hansen et al., 2013; Osterbye et al., 2020; Malovichko and Zhu, 2017), it is now possible to display complete proteomes in a single slide (Lagatie et al., 2017; Yan et al., 2019), opening the door to high-throughput serological screenings.

Chagas Disease, also known as American trypanosomiasis, is a lifelong infection caused by the protozoan parasite *Trypanosoma cruzi*. Despite being discovered ∼100 years ago, the condition remains a major social and public health problem in Latin America and is regarded as a neglected tropical disease by the World Health Organization (Pérez-Molina and Molina, 2018).

After an initial infection, the parasite evades immune mediated elimination and mounts long-lasting chronic intracellular infections. Due to the low parasitemia observed during the chronic stage of the disease, serological methods are the preferred choice for diagnosis of infection. Although available diagnostic tests give satisfactory results in most cases, there is currently no reference (“gold”) standard for diagnosis of infection hence discordant results remain a possible cause of undetected cases (Balouz et al., 2017; Guzmán-Gómez et al., 2015; Moure et al., 2016). Also, there are urgent needs to improve or fill vacant niches with customized serological tools and assays to monitor existing treatments or clinical trials (Granjon et al., 2016; Jurado Medina et al., 2021; World Health Organization, 2012) and to detect the early onset of Chagas Disease pathology (Pereira Nunes et al., 2018), both in active case finding and management and in epidemiological and disease surveillance programmes (Peeling, 2015; Peeling et al., 2017).

In this article, we report on the comprehensive survey and characterization of the human antibody responses and specificities against *T. cruzi* using state of the art high-density peptide arrays. This survey investigated the antigenicity of the predicted proteomes of two *T. cruzi* strains in 71 Chagas Disease subjects from diverse human populations across the Americas. As a result, we produced a comprehensive atlas of antigens and their epitopes, providing a unique resource for understanding adaptive immune responses against this parasite and to devise improved serological immunoassays for tackling Chagas Disease.

## RESULTS

### CHAGASTOPE-v1: design of a high-density peptide array for antigen and epitope discovery

We used high-density peptide arrays to perform a high-resolution antigen discovery screening and epitope mapping for complete *T. cruzi* proteomes. We designed an array that included protein sequences encoded in the genomes of two *T. cruzi* strains: the genome reference CL Brener strain (19,668 proteins, Discrete Typing Unit (DTU) TcVI, hybrid) (El-Sayed et al., 2005), and the Sylvio X10 strain (10,832 proteins, DTU TcI, non-hybrid) (Franzén et al., 2012). The selection criteria considered the epidemiological relevance of these representative lineages in the context of Chagas Disease, and the fact that TcVI strains are hybrids of ancestral DTUs TcII and TcIII (Brenière et al., 2016; Zingales, 2018), resulting in most of its genes being represented in the genome by their two ancestral allelic versions (El-Sayed et al., 2005; Souza et al., 2011). Therefore, by using strains from DTUs TcI and TcVI we maximized the display of relevant peptide variants with only two genomes.

Based on this analysis and the high-density peptide array capacity, we produced a tiling display of 30,500 *T. cruzi* proteins using 16mer peptides with an offset of 4 amino acids (overlap of 12 residues between consecutive peptides). The resulting array containing 2,441,908 unique peptides was named CHAGASTOPE-v1 and was used for the discovery screening (see Figure 1). Additional details on the contents of the CHAGASTOPE-v1 array design are available in the Methods and in the Supplementary Tables S1, S2, S3 and S10.

**Figure 1.**
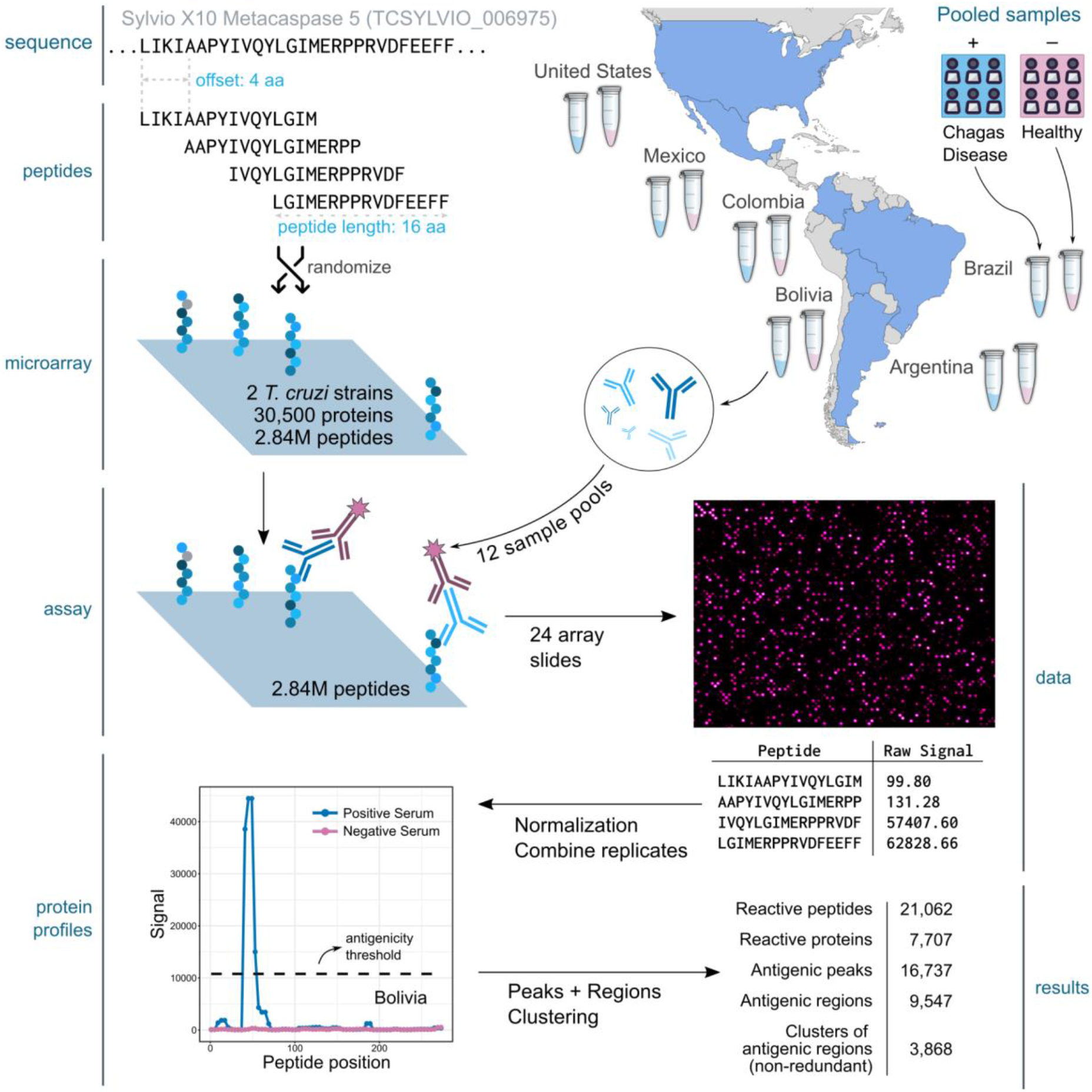
Summary of the Discovery Screening. The figure shows a schematic representation of the steps followed to analyze two T. cruzi proteomes (CL-Brener and Sylvio X10) using pooled serum samples across the Americas (one pool from infected individuals and one from healthy subjects from Argentina, Brazil, Bolivia, Colombia, Mexico and the United States). The protein used for this example is the metacaspase 5 protein from Sylvio X10 (TCSYLVIO_006975) which has 291 residues and was represented in each array using 69 peptides (16mers with an offset of 4 aa), of which only the first 4 peptides are shown.

### Discovery Screening: A high-content serological screening reveals diverse antibody repertoires in Chagas Disease and novel antigens

Using the CHAGASTOPE-v1 array design, we performed a discovery screening using pooled serum samples from positive donors and paired negative sample pools from healthy subjects for 6 different geographical regions across the Americas (Figure 1). In total, we profiled 12 different pooled serum samples, 6 from infected subjects and 6 from healthy subjects (see Supplementary Table S2 for details). To test for cross-reacting epitopes, two additional pools were profiled: a pool from leishmaniasis-positive individuals and a matched pool of leishmaniasis-negative (also Chagas-negative) samples from the same geographic area. All 14 samples were assayed in duplicate, and all technical replicates had high signal correlation (see Supplementary Figure S2).

After normalizing the antibody-binding fluorescence signal across experiments, we reconstructed each original protein sequence, used consecutive peptides to perform signal smoothing (to remove outliers, see Methods) and generated visualizations of the antibody-binding signal for each protein and for each assayed sample (see example in Figure 2a and the full list in Supplementary File S2).

**Figure 2.**
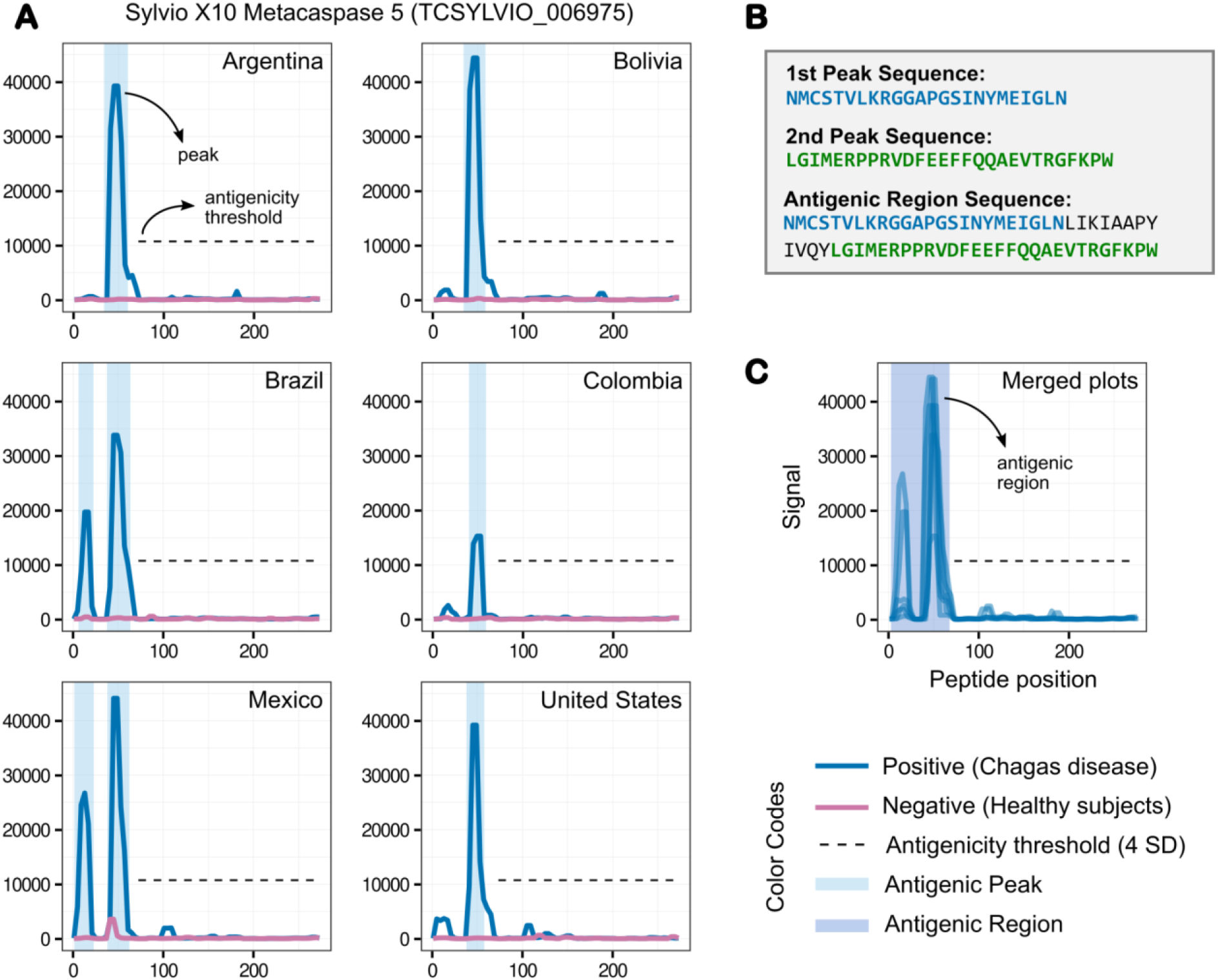
Antibody-binding profiles, peaks, and regions. The normalized fluorescence signal of each peptide in any given protein was used to produce the antibody-binding profiles for an antigen. The y axis shows fluorescence units. The x axis shows peptide positions along the protein sequence. Each subplot was produced using data from 4 high-density peptide arrays (2 replicas for Chagas disease subjects and 2 for matched healthy subjects, see main text). **A)** Reactivity subplots for different sera pools for the Sylvio X10 metacaspase 5 protein. The antibody-binding profiles are shown in blue for the chagas-positive sample pools (infected) and in magenta for the chagas-negative pools (healthy). We used a conservative 4SD antigenicity threshold throughout the discovery phase, which can be seen as a black dashed line. **B)** Peptide sequences of the reactive peaks and regions in A. **C)** Reactivity plot merging data from all sample pools. The figure serves to illustrate how we defined peaks (groups of consecutive peptides over the signal threshold) and antigenic regions (groups of neighboring peaks).

To identify the more reactive proteins, we compared signals obtained across experiments with pools of Chagas-negative and Chagas-positive subjects and defined an antigenicity signal threshold. We chose a very conservative threshold of 10,784.80 fluorescence units (the statistical mode plus 4 standard deviations). Any group of two or more consecutive peptides above this threshold was defined as an antibody-binding peak (see Figure 2), resulting in 18,199 peaks for the Chagas-positive subjects. We also observed 3,644 reactive peaks in Chagas-negative subjects (see Supplementary Figure S4). After removing these cross-reactive peaks we obtained 16,737 Chagas-specific antigenic peaks across 7,707 proteins.

Because some peaks were either close or partially overlapping with one another (in the same analyzed sample or across different samples, see Figure 2c), we combined neighboring peaks into non-overlapping antigenic regions. This resulted in 9,547 antigenic regions across both proteomes (see Methods). Furthermore, because the analyzed *T. cruzi* genomes have several large gene families, a significant number of reactive regions displayed evident sequence similarity amongst them. Hence, we grouped these antigenic regions into 3,868 non-redundant clusters based on sequence similarity using protein BLAST (see Methods). The identification of reactive peptides, peaks and regions, and their cognate antigenic proteins is summarized in Table 1.

**Table 1.** Discovery screening finding summary. This table displays the number of reactive peptides, peaks, regions and proteins found in our first screening. When analyzing the number of peptides each unique sequence was only counted once.

|  | CL-Brener | Sylvio X10 | Pangenome |
| --- | --- | --- | --- |
| **Total Proteins** | 19,668 | 10,832 | 30,500 |
| **Total Peptides (non-redundant)** | 1,809,351 | 1,187,499 | 2,441,908 |
| **Reactive Peptides (non-redundant)** | 16,435 | 7,709 | 21,062 |
| **Antigenic Peaks** | 12,720 | 4,017 | 16,737 |
| **Antigenic Regions** | 6,710 | 2,837 | 9,547 |
| **Reactive Proteins** | 5,328 | 2,379 | 7,707 |
| **Clusters of Antigenic Regions (non-redundant)** | - | - | 3,868 |

### The immune responses in Chagas disease subjects are highly diverse

The complete map of measured antibody-binding reactivities across pooled samples provided a broad view of the diversity of the antibody repertoire developed in response to *T. cruzi* infections. We next analyzed how reactive regions were shared amongst pooled samples and observed a large set of non-shared reactive epitopes in each sample (see Figure 3). These were not technical artifacts as they were reproducibly identified in the technical replicates and were also supported by the reactivity of overlapping neighboring peptides (pseudo-replicates within each experiment).

**Figure 3.**
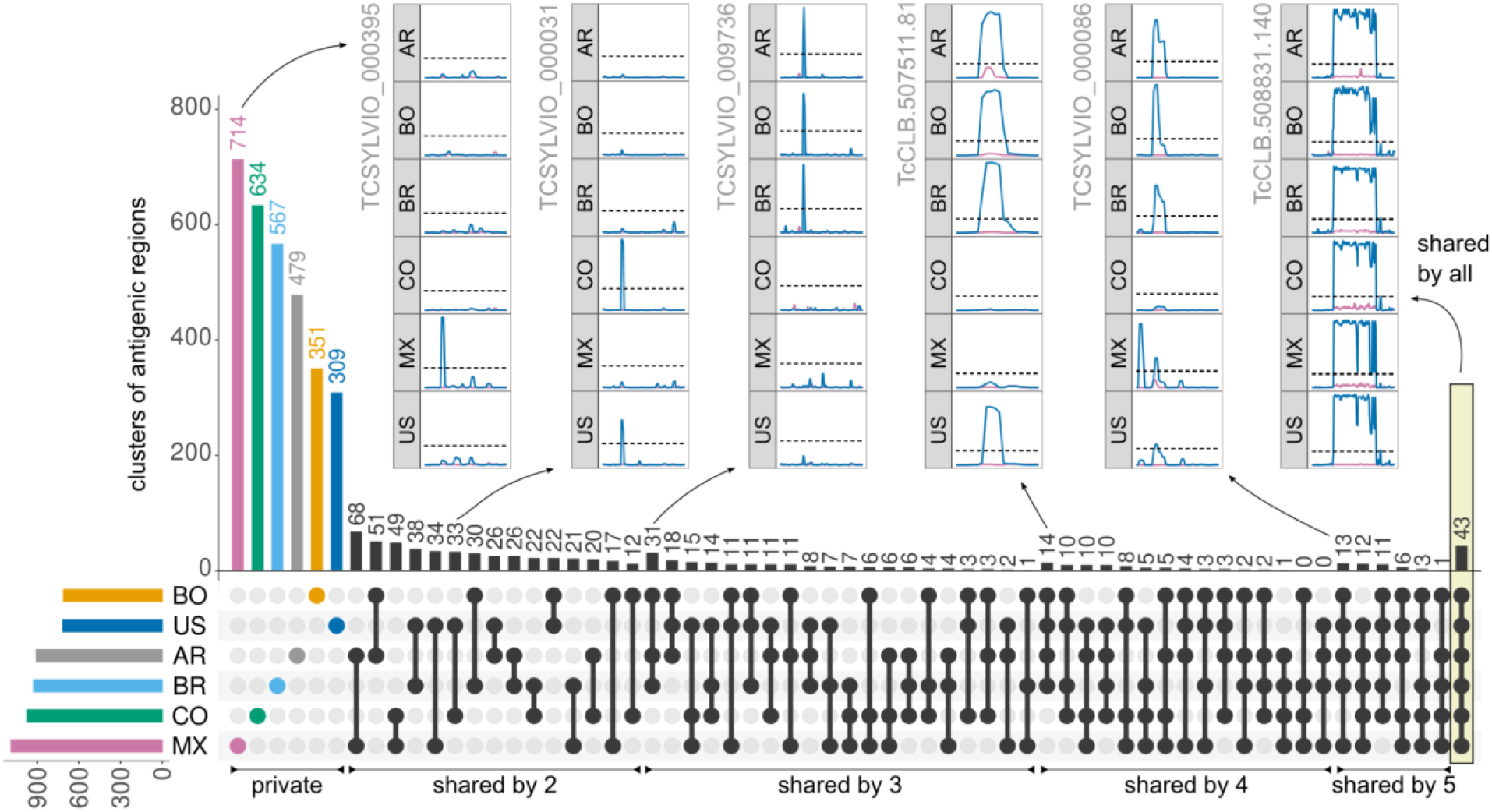
Diversity of T. cruzi antibody-specific responses in pooled serums. Non-redundant clusters of reactive regions in the analyzed T. cruzi proteomes (clustered by sequence similarity) were counted on all intersections of the 6 analyzed pooled samples. A cluster was antigenic in each sera pool when at least one of its regions was antigenic in that pool. The UpSet Plot (Lex et al., 2014) displays a histogram with these counts (top), as well as a visual depiction of all the set intersections (bottom, black). The colored histogram at the bottom left shows the counts of total reactive regions. Pooled samples are AR = Argentina; BR = Brazil; BO = Bolivia; CO = Colombia; MX = Mexico; US = United States. The insets show antibody-binding profiles (as in Figure 2) for several selected antigens.

This set of non-shared (private for pool) antigenic regions was followed by a long tail of shared epitopes with increasing seroprevalence. This observation suggests that the antibody response in Chagas disease is derived from a large and diverse set of antibody-producing B-cell clones. Particularly important for serology-based applications are the large set of shared clusters of antigenic regions across Chagas disease subjects (166 shared by at least 4 of the analyzed pooled samples), including 43 clusters that were reactive in all samples (see Table 2, and Supplementary Table S6). The full list of antigenic regions for all clusters can be found in Supplementary Table S12.

**Table 2.**
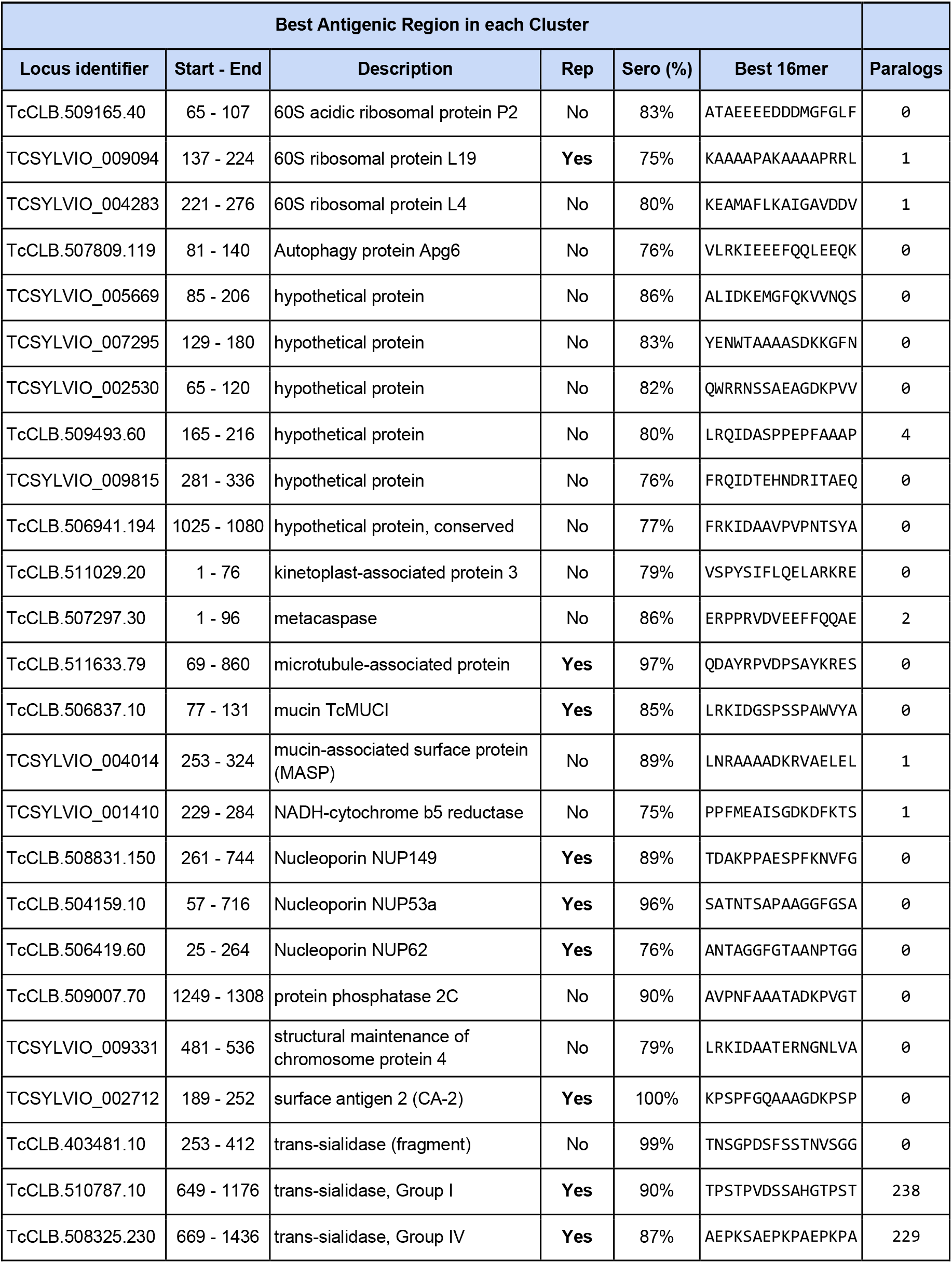
Antigens reactive in all Chagas-positive pooled serum samples. A selection of 20 clusters of antigenic regions from those showing reactivity against all assayed positive pooled samples. The clusters were selected based on cross-reactivity, average fluorescence signal and protein family. We excluded those clusters that showed cross-reactivity against Leishmaniasis pooled samples. For each cluster, we show a representative antigenic region with the highest seroprevalence for that cluster. Rep = repetitive (contains internal tandem repeats); Paralogs = number of additional non-allelic copies of the gene.

We also screened the same array design against a pool of samples from Leishmaniasis-positive individuals to identify cross-reacting epitopes. Overall, there was very low cross-reactivity of *T. cruzi* peptides against this pool. Out of the 3,868 clusters of antigenic regions, only 104 (2.7%) had reactivity against this leishmaniasis sample (using the same threshold used for *T. cruzi* samples). This included 5 of the 43 clusters that had reactivity in all Chagas-positive samples, although for 3 of these only a small percentage of the regions in the cluster were cross-reactive.

### Individual patient resolution provides insights into the diagnostic value of identified antigens and epitopes

The previous screening provided ample information on the diversity and specificities of the antibody repertoire of Chagas disease patients. However, the use of pooled samples limited the analysis of the data. Next, we increased the epitope mapping resolution and assessed the reactivity of epitopes for individual patient samples.

Because 99% of peptides showed no antibody-binding at the screening stage, we removed these from the new design to focus only on the antigenic regions. Working with these smaller regions of the proteome allowed us to increase the epitope mapping resolution, using 16mers with an overlap of 15 residues between consecutive peptides.

This second peptide design (named CHAGASTOPE-v2) included peptides from the 9,547 protein regions that were both antigenic (reactive with Chagas-positive samples) and that showed no signal from healthy control subjects. To ensure that entire reactive peaks in each region were included in the design, we included up to 32 additional peptides from the surrounding non-reactive borders of each region (16 from each side). This contributed to the v2 array design with 241,772 unique peptides from this set.

To expand resolution of epitopes down to individual patients, we used sectorized high-density peptide arrays to assay more samples (primary antibodies) in parallel in the same slide (see Supplementary Table S1). In these 12-plex arrays, the same set of CHAGASTOPE-v2 peptides were replicated in each sector. A total of 12 CHAGASTOPE-v2 arrays (144 sectors) were used to analyze 71 individual serum samples in duplicate, 33 of which were from the pools analyzed in the previous step. A set of 38 additional serum samples were analyzed from other Chagas-positive subjects from the same 6 geographic regions (see Supplementary Table S2). The replicas showed high Pearson correlation coefficients (>0.8, see Supplementary Figure S3). Because these arrays have a much lower content of non-reactive peptides, we derived the antigenicity threshold independently resulting in a threshold of 5,814.81 relative fluorescence units (statistical mode plus 2.4 standard deviations, see Methods).

Using these arrays, we obtained a fine look at antibody-binding regions in proteins (see Figure 4) and extracted detailed seroprevalence information at the level of proteins and individual epitopes. As shown in the examples in Figure 4, the resolution of antibody-binding reactivity in each individual provides information on the seroprevalence of each antigenic region. A novel antigen with a single antigenic region is shown in Figure 4a (reticulon-domain containing protein, TCSYLVIO_003288). The observed seroprevalence at the level of the whole protein is aligned with the seroprevalence observed at the single-epitope level. A different extreme example is the hypothetical protein encoded by gene TCSYLVIO_005669, another novel antigen (see Figure 4B). This protein displayed a contiguous antigenic region composed of 7 different antibody-binding peaks (epitopes), each with a unique set of signal and seroprevalence characteristics. In this instance the global seroprevalence for this antigen was not aligned with the seroprevalence of individual epitopes. Finally, for known *T. cruzi* antigens, resolution of how antibodies from different individuals recognize these antigens provide insights to improve diagnostic reagents. The antibody binding profiles of the Ag36/MAP antigen (Ibañez et al., 1988) for US samples are shown in Figure 4c. As shown in the figure, the antigenic repetitive unit is not recognized in the same way by all individuals. While most individuals display reactivity for all overlapped epitopes in this repeat, some individuals (e.g., US_P3, US_E2) have antibodies with different binding preferences along this antigenic repetitive unit.

**Figure 4.**
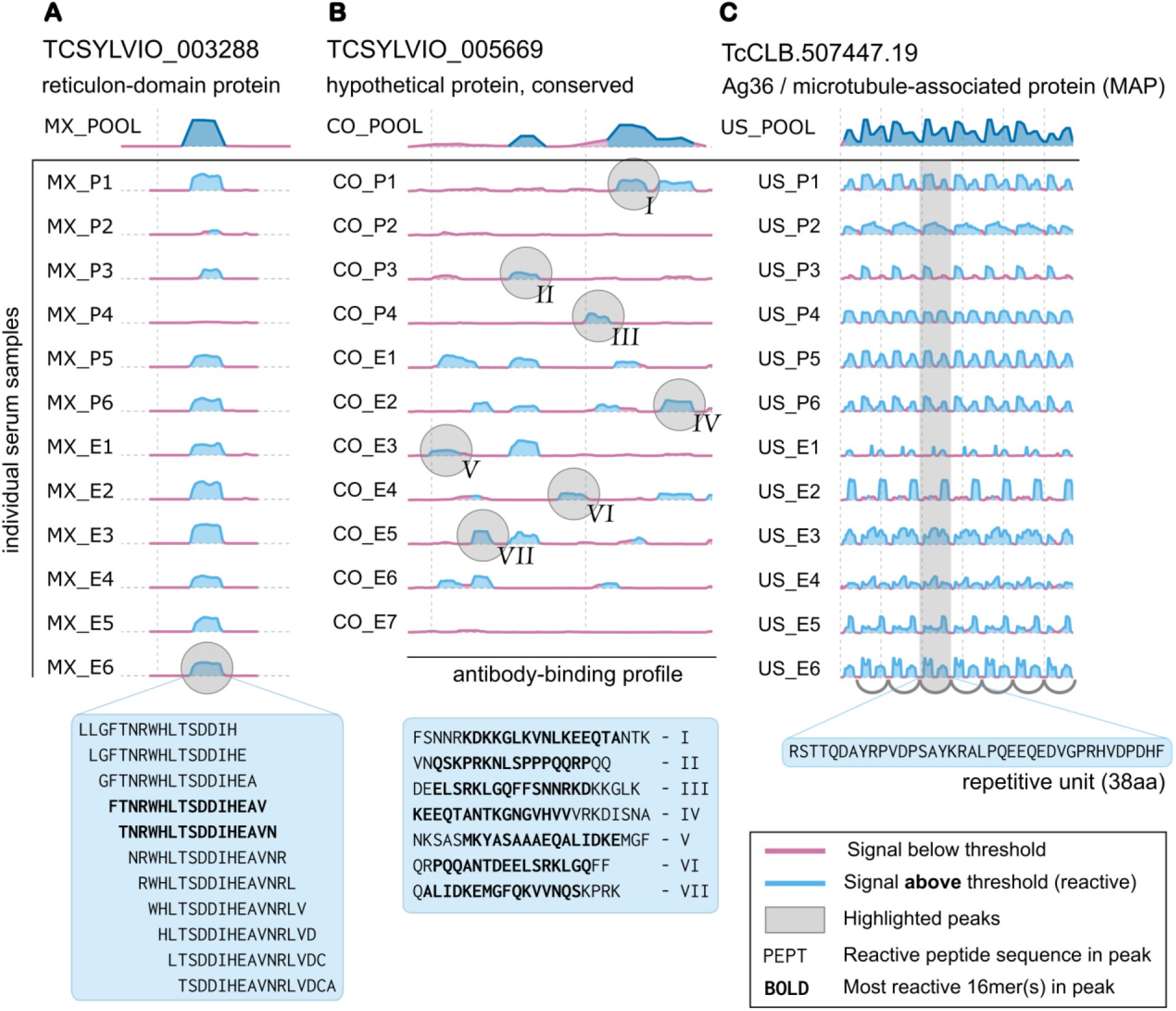
Individual patient resolution and epitope mapping of Chagas Disease antigens. Example antibody-binding profiles obtained using CHAGASTOPE-v2 arrays. In all cases the reactivity of the pooled samples in the CHAGASTOPE-v1 discovery screening is shown at the top (dark blue). See Supplementary Table S2 for the codes of patient serum samples (MX = Mexico, CO = Colombia, US = USA). **A)** example antigen with a single reactive region, showcasing peptides with signal above threshold (and most reactive peptides in **bold**). **B)** non-repetitive antigen displaying multiple reactive peaks (marked using **roman numerals**), and the corresponding peak sequences below. **C)** repetitive antigen displaying heterogeneous recognition of the repetitive unit by different Chagas-positive individuals.

Supplementary Files S3 and S4 contain the antibody-binding profiles for all antigens and all analyzed patient samples tested in CHAGSTOPE-v2 (plots similar to those in Figure 4). This is a rich dataset for future studies on the serology and immune responses of Chagas disease patients and serves as a reference for other infectious diseases.

Resolution of antibody-binding to defined peptides in each subject allowed us to explore reactivity against *T. cruzi* strains analyzed in this study. Figure 5 summarizes the reactivity of 33 subjects against peptides exclusive to CL-Brener or to Sylvio X10. Subjects displayed reactivity to an average of 4,088 CL-Brener peptides (min: 3,174; max: 5,113, std: 404) and 955 Sylvio X10 peptides (min: 712; max: 1,249; std: 151). Analysis was performed using z-scores because of the difference in numbers of peptides in each set (CL-Brener is a larger hybrid genome, see Methods). Most samples from Mexico displayed higher relative reactivity against peptides from the TcI / Sylvio genome (as well as subjects AR_P1, CO_P1, and US_P1). Other subjects, particularly from Argentina, Brazil and Bolivia showed higher relative reactivity against TcVI / CL-Brener (e.g. AR_P2, AR_P4, BO_P3, BR_P4). While the design of these arrays prevents a more detailed serotyping analysis, these serological reactivity signatures suggest differences in the infecting *T. cruzi* lineages in these subjects (the complete analysis with all serum samples is available in Supplementary Figure S5).

**Figure 5.**
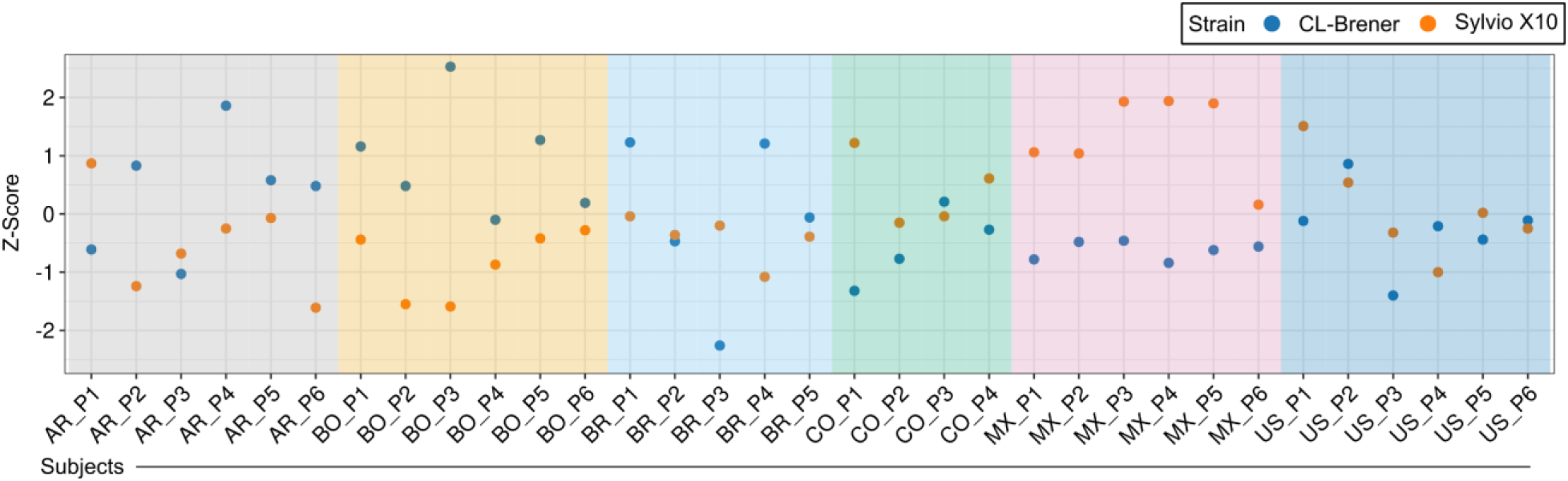
Peptide antigenicity of individual patient samples against T. cruzi strains. Counts of unique reactive peptides in the two analyzed T. cruzi strains were standardized using two sets of z-scores, one per strain (see Methods). A pair of Z-scores for each subject informs the relative reactivity observed.

### Diversity of individual immune responses, revisited

In the discovery screening we observed that most antigenic regions were private, meaning that they were reactive in only one pooled sample. Here, we revisited this analysis using data from the CHAGASTOPE-v2 arrays. We analyzed the 3,868 clusters of antigenic regions identified at the discovery stage (3,054 private/non-shared, 814 shared) now using 71 individual serum samples, and confirmed our observation of a diverse and quantitatively large set of private (non-shared) immune responses (Figure 6).

**Figure 6.**
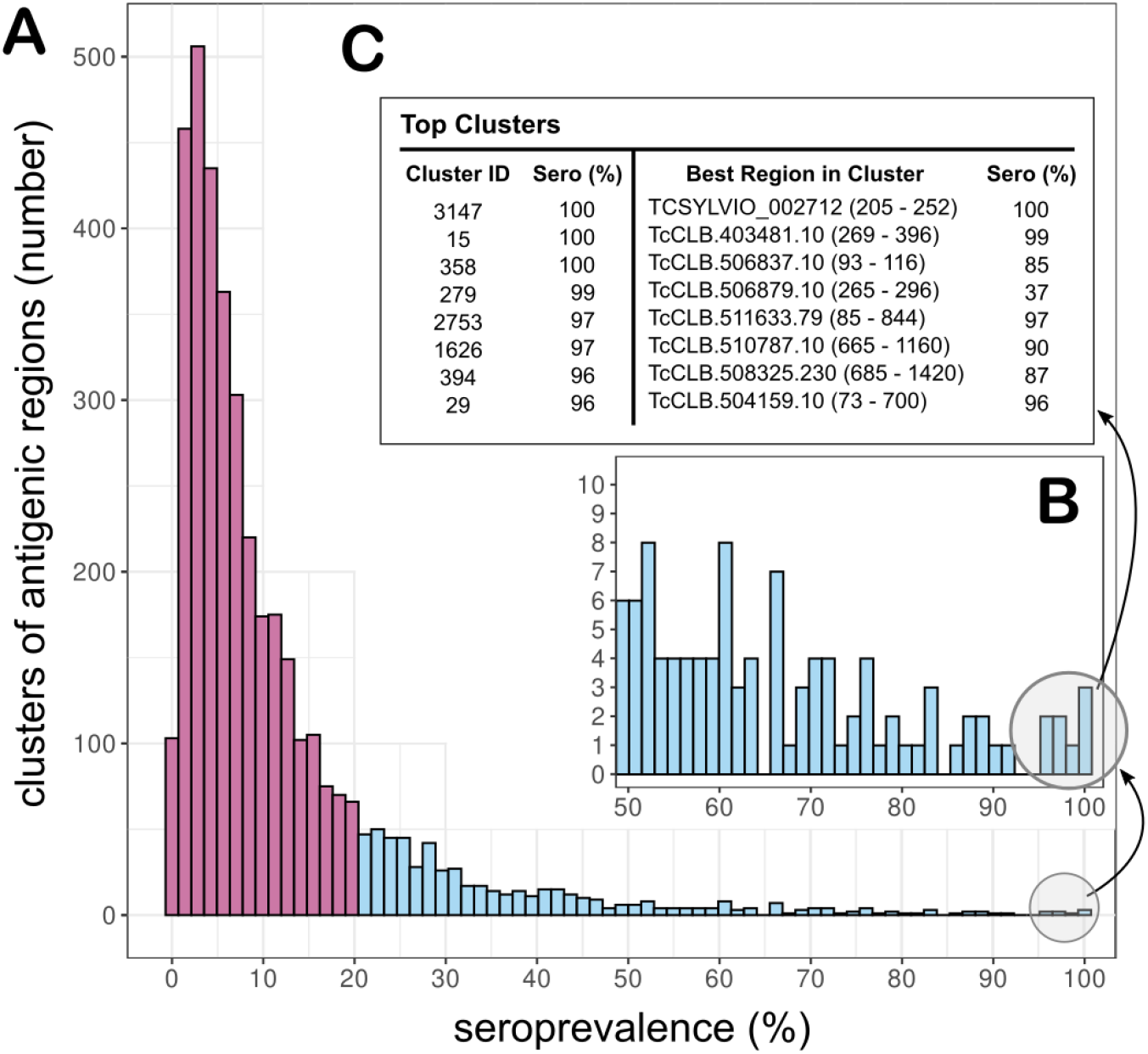
Profiling of individual antibody responses reveals the diversity of anti-T. cruzi specific antibodies. Non-redundant clusters of reactive regions (clustered by sequence similarity) were counted on all intersections of the 71 analyzed individual samples. **A)** histogram showing the number of clusters that were reactive in each fraction of subjects (shown as seroprevalence). In magenta: clusters reactive in 20% or less of the individuals (mostly private, non-shared). **B)** Inset zooming in on clusters reactive in at least 50% of the subjects. **C)** Table with details on 8 clusters that were reactive in most individual serums. The regions shown for each cluster have the highest seroprevalence among the regions in that cluster.

Most clusters of antigenic regions were detected by a low number of individuals, with 65% of the clusters showing a seroprevalence of 10% or less, and 85% displaying a seroprevalence lower than 20% (magenta in Figure 6a). When analyzing these 71 samples individually, we observed 88,236 reactive peptides (non-redundant); however, each individual displayed reactivity to 5,841 peptides on average, which is in agreement with the observed large repertoire of private antigens.

Table 3 provides additional information on the clusters with higher seroprevalence, highlighting 25 clusters with regions detected by at least 75% of the subjects. While some of these regions were already known *T. cruzi* antigenic proteins, there were novel ones identified, which are of interest for improving diagnosis and treatment of Chagas disease. Information for all clusters can be found in Supplementary Table S7.

**Table 3.**
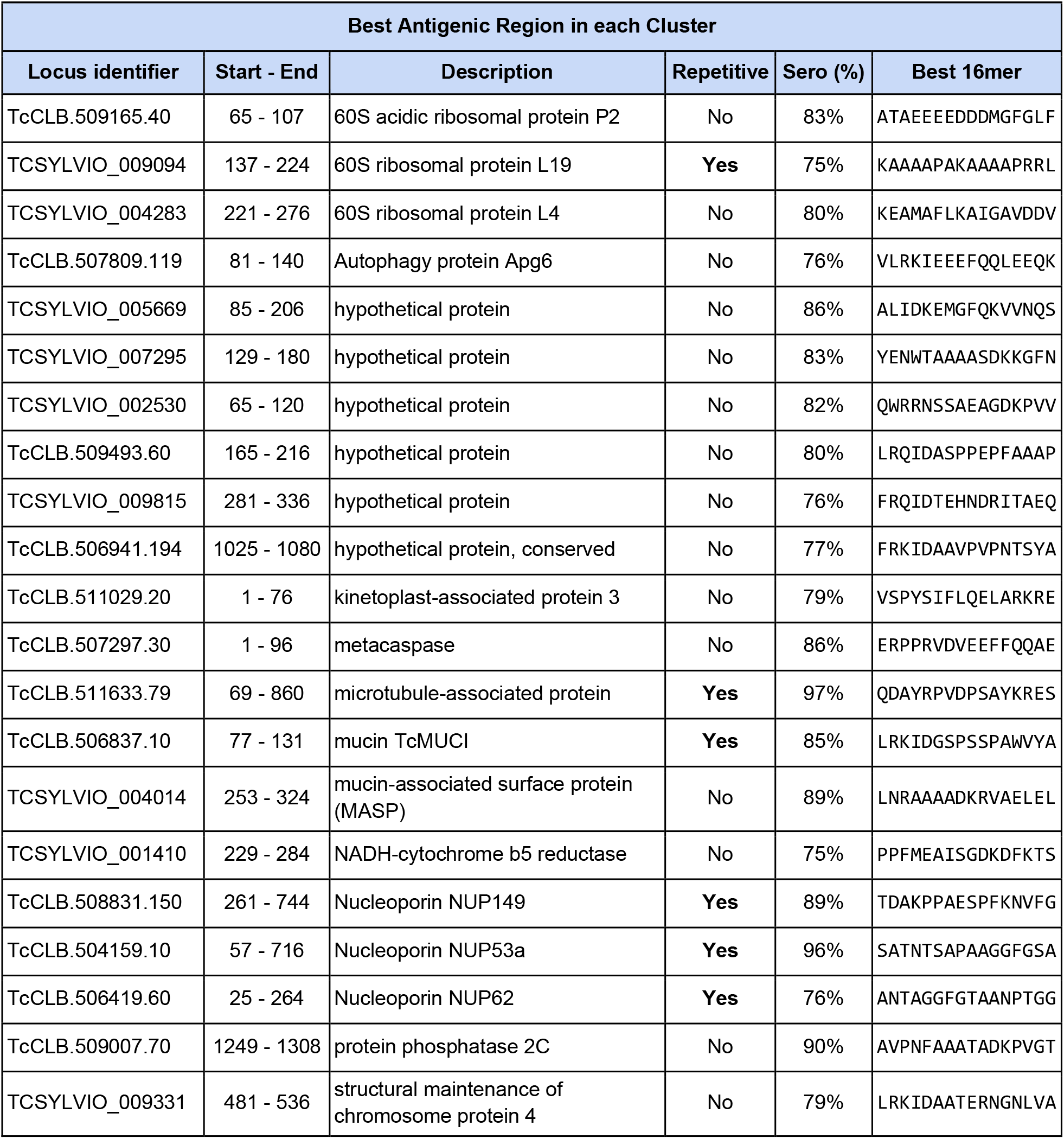
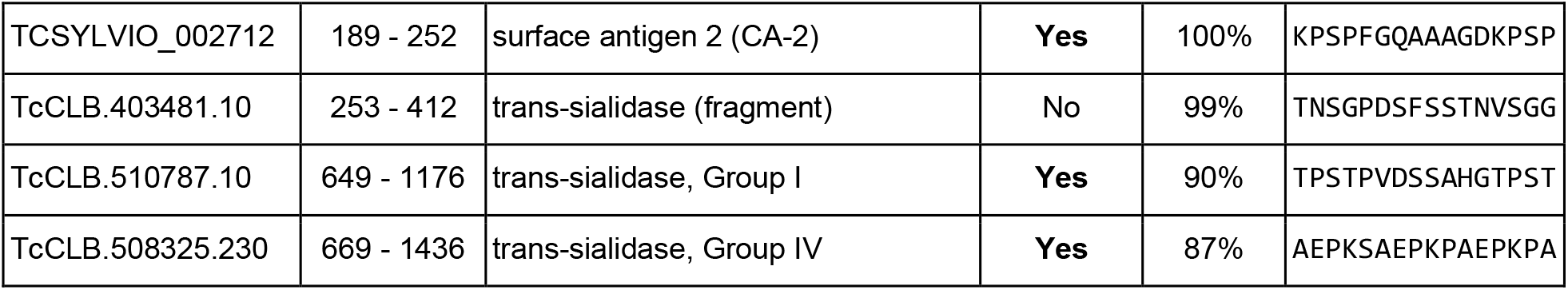
Antigens reactive in most Chagas-positive individual serum samples. A selection of most seroprevalent clusters of antigenic regions where both the cluster and at least one of its regions was detected by 75% of the 71 individual serum samples. For each selected cluster, we show a representative antigenic region with the highest seroprevalence for that cluster (“Sero” indicates the restricted seroprevalence for the selected protein region).

### Most human immune responses converge on the same fine epitope specificities

To investigate the modes of recognition of antigens and epitopes by different individuals we performed alanine-scan mutagenesis of selected epitopes. Because substitution with alanine removes all side chain atoms past the β-carbon, the effects of individual alanine mutations can be used to infer the roles of individual side chains. In the CHAGASTOPE-v2 array design we included mutants designed *in silico* for 232 selected antigenic peaks found in the discovery screening. For each selected peak, the best 20mer sequence was used as a guide to probe antibody-binding by replacing each residue for an Alanine (except when the residue itself was Alanine, in which case we replaced it by Glycine), see Figure 7a. This was done for all 5 overlapped peptides (16mers) in each antigenic sequence; hence the same mutation was assayed several times, in different positions. Supplementary Figure S6 and Methods provide a full description of the procedure.

**Figure 7.**
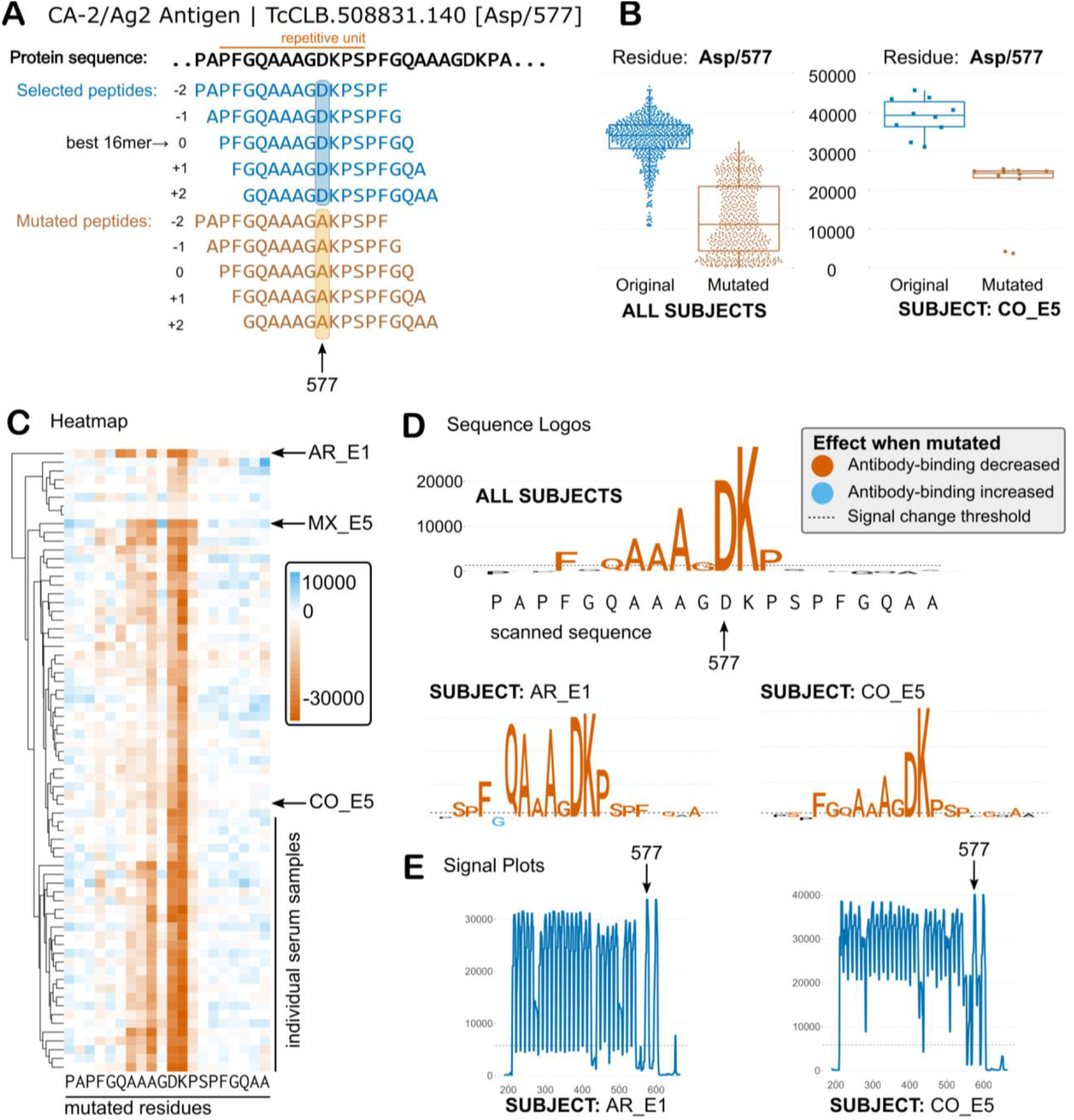
Single-residue scanning mutagenesis of T. cruzi antigens. **A)** Schematic representations of single-residue mutagenesis for one repetitive antigen (only 1 residue is mutated in this depiction for clarity). **B)** Average signal of the original and mutated peptides for the Asp/577 residue for all positive sera (left) and for one selected subject (right). **C)** Heatmap showing the effect of mutations on antibody binding (signal change from original to mutated peptides) for all residues and for all sera. Mutations that decrease antibody-binding are shown in different shades of orange, while those that increase binding are shown in shades of blue. Columns = residue positions, Rows = individual serum samples. **D)** Sequence logos summarize data for all positive sera (Core residues), or for individual cases (y-axis: signal change in fluorescence units). Colors follow heatmap. **E)** Antibody binding signal plots for selected subjects.

When performing single-residue mutational scanning over different epitopes, we observed that antibody-binding was diminished consistently for some amino acid residues. These key residues define the ***core functional*** residues of each epitope, which represent residues likely in contact with the paratope in the Fab (antibody-binding) region of the immunoglobulins. As seen in Figures 7 and 8, the core residues were approximately 5 to 9 residues for these epitopes (9 was the most frequent value, with 65% of the epitopes having between 7 and 11 ***core*** residues, see Supplementary Fig S11). Interestingly these core residues were mostly shared across positive sera (see heatmaps in Figure 7, and Supplementary File S7), and hence suggest that immune responses from different individuals converge on similar modes of binding.

**Figure 8.**
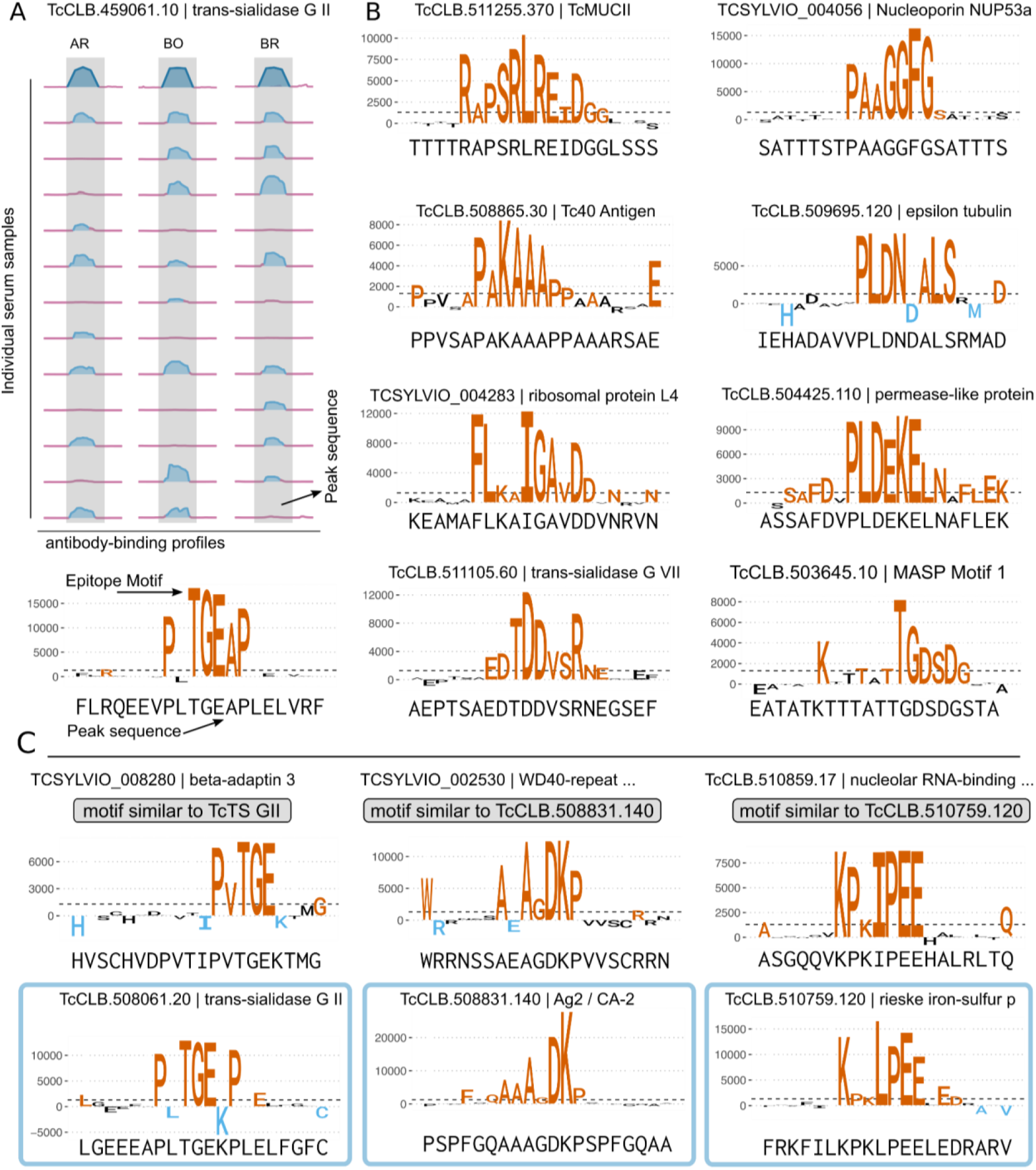
A gallery of functional epitope motifs. **A)** Antibody-binding profiles for one selected antigen, similar to Figure 4 (only a subset of 36 of the 71 analyzed sera is shown for clarity). The best 20mer (peak sequence) was subjected to single-residue mutagenesis (see Methods), and the results were summarized as sequence logos. **B)** Sequence logos showing epitope motifs for additional antigens. Colors for residues that affect antibody-binding are similar as in Figure 7. **C)** Example cases of antigens with similar core residues (candidate mimotope motifs). Those with higher signal and/or seroprevalence are boxed.

As an example, the Ag2/CA-2 repetitive *T. cruzi* antigen (Buschiazzo et al., 1992), one of the most seroprevalent antigens in our screening, has a functional core defined by the AAGDK residues of the 12mer repetitive unit (see Figure 7). Even individuals that showed low signal in the arrays or were borderline negative for this antigen with our signal threshold displayed the same residue fingerprint characteristic of this epitope (see Supplementary Figure S7). This suggests that this convergent specificity on the antibody-binding mode is maintained across high-and low-titer responses.

Despite overall convergence on key residues, we also observed differences in the way that antibodies from some individuals bind to epitopes. For Ag2 the individual AR_E1 displays higher relevance for residues 572Q and 573A (see Figures 7d and 7e) even when the signal profile of the protein was similar (see Figure 7f). Also, the high resolution of this technique allowed us to observe additional residues that in some individuals produce unique effects on antibody-binding (when mutated). For example, at least 7 individuals showed increased antibody-binding when 576G was mutated to an Alanine (with individual MX_E5 displaying the most extreme effect see Figure 7c). Examples of other cases of differential epitope recognition by individual subjects are presented in Supplementary Figures S9 and S10.

Similar observations can be made for other antigens. In the case of the Gim5A protein, the core epitope is defined by 4 core residues with a very high mutational signal (LPED), and 3-4 additional residues with more heterogeneity in the way that antibodies from different individuals recognize this epitope. For example, the 136K-138A residues preceding the core epitope motif were relevant for antibody binding for ∼half of the positive serum samples for this antigen (see Supplementary Figure S8), whereas these residues were not relevant for the antibodies developed by other subjects (e.g., BO_E6 and AR_E1).

The high resolution achieved by this strategy also allowed us to reveal cross-reactivity between epitopes that bear low sequence similarity. These putative antigens showed no detectable similarity to other antigens (hence they were not clustered together at the BLASTP stage) but displayed highly similar (or identical) key residues in this mutational analysis. One such example was a hypothetical protein (TCSYLVIO_002530) for which mutagenesis scanning revealed the characteristic AGDK core of the Ag2/CA-2 antigen (see Figure 8C and Supplementary File S7). There is almost no similarity between the protein sequence of Ag2 and this hypothetical protein, and even when looking at the scanned 20mer, there are only 5/6 shared residues. The fact that these residues constitute the core functional residues of the Ag2/CA-2 antigenic repeat suggests that the TCSYLVIO_002530 protein is not a *bona fide* antigen and the reactive region detected in our screenings is instead a mimotope.

Examples of other candidate mimotopes are shown in Figure 8C: the beta-adaptin 3 protein (TCSYLVIO_008280) with a core functional epitope highly similar to that of the trans-sialidase protein (TcCLB.459061.10), and the nucleolar RNA-binding protein (TcCLB.510859.17) with a functional core similar to the rieske iron sulfur protein (TcCLB.510759.120), amongst others (see also Supplementary File S7). In all these cases, the normalized signal observed in the arrays for the true positive antigens was ∼2 fold higher than the signal observed for the candidate cross-reactive proteins.

A detailed account of all the core epitope motifs revealed by this study will likely be further explored by researchers in the field. Supplementary Tables S8 and S9 provide a summary of the key residues in other epitopes analyzed, and similar visualizations of these data are provided for all mutagenized epitopes as Supplementary File S7.

## DISCUSSION

We have produced a detailed characterization of the antibody specificities against linear epitopes in patients with Chagas disease, described in depth here for the first time. Previous studies have shown that some pathogens deliberately induce short-lived, polyclonal plasma cells to dilute long-lived, specific antibody responses (Nothelfer et al., 2015). In contrast, *Trypanosoma cruzi* induces a massive clonal expansion of B-cells during the acute phase leading to production of parasite-specific and autoreactive antibodies as well as high levels of antibodies with unknown specificity (Bermejo et al., 2011; Minoprio, 2001). Our description of a large and diverse number of specificities, composed of mostly non-shared epitopes (low seroprevalence at the population level), support these previous observations, and provide information on the targets of these antibodies.

The power of these massively parallel serological assays lies in the delineation of the responses of individual patients to each identified epitope, hence producing a rich human seroprevalence matrix for these antigens. Our analysis uncovered >3,800 non-redundant antigenic regions, both public (shared) Chagas antigens as well as private individual anti-*T. cruzi* responses, in 71 individuals from different human populations across the Americas.

A consequence of the experimental platform we used to detect antibody-binding, is the inherent bias towards linear epitopes (likely missing most conformational epitopes). This is evident in our failure to detect antibody binding to at least one known *bona fide* antigen: Ag1 (Ibañez et al., 1988), also known as FRA or JL7 (da Silveira et al., 2001), a cysteine protease (clan CA, family C2, CL-Brener Locus ID: TcCLB.505985.9) which is a component of commercial kits for the diagnosis of *T. cruzi* infection. No reactivity was observed in our short peptide screenings, suggesting that the epitope(s) in this antigen may be conformational.

To identify cross-reactive epitopes against a co-endemic disease caused by a related trypanosomatid parasite, we also assayed a pool of leishmaniasis-positive serum samples against *T. cruzi* peptides from complete proteomes. This is important, as many Chagas disease false positives are suspected to be a consequence of Leishmania spp. cross-reactivity (Abras et al., 2016). We identified 888 *T. cruzi* peptides that were cross-reactive with these leishmaniasis samples (data in Supplementary Table S4). While the number of leishmaniasis samples in the pool may be small (n = 6), the ability to screen at this scale outweighs this potential limitation.

Mutational scanning of a large epitope set revealed key residues for antibody-binding. The precision of this type of analysis led to identification of cases where antibodies from different subjects converged to a shared mode of antibody-recognition for the studied epitope, and cases where the same epitope had divergent (alternative) modes of recognition by different individuals (see Figures 7, Supplementary Figures S9 and S10, and Supplementary File S7). Both convergence and divergence in B cell clonal lineages has been observed (Krause et al., 2011) leading to recurring motifs for the recognition of viral protein antigens (Xu et al., 2013). This mutagenesis data represents a rich resource to guide downstream applications. Exploration of these data in the context of antigen variants identified from genomic sequencing of *T. cruzi* isolates may improve assay reagents.

To our knowledge, this Atlas is the largest collection of Chagas disease antigens and epitopes described to date, and the first dataset providing fine resolution of seroprevalence to epitopes in humans. Due to the breadth and diversity of the clinical samples analyzed, this study also provides a large set of experimentally validated negative data (non-antigenic proteins and peptides). This is almost always overlooked, but it represents a highly valuable dataset for training of predictors, which often need to work under the assumption that proteins with no previous information on their antigenicity are non-antigenic (Larsen et al., 2006; Ricci et al., 2021). The datasets from the primary discovery screening also provide a large corpus of data on dominant *T. cruzi* peptides reactive to sera from healthy subjects from different human populations.

The observations and the data produced in this study reflect a snapshot in time of the antibody repertoires of each subject. Many questions about these repertoires remain. What is the nature of the private (non-shared) set of antibody specificities? Which epitopes are the targets of short-lived responses? And which are the targets of long-lived responses? The observed low seroprevalence of a large fraction of antigens (non-shared responses) may be explained if this is a fluctuating repertoire. It is thus tempting to speculate that private antigens (as described in this work) may be the target of short-lived or weak antibody responses. Under this scenario, the B-cell clones producing these antibodies may decay after some time, and thus the observed feature of being unique to one or very few individuals in these snapshots may be the telltale of these short-lived immune responses. This agrees with the current view of the complex and focal dynamics of Chagas disease in the host (Lewis and Kelly, 2016; Pérez-Mazliah et al., 2021; Ward et al., 2020), where waves of parasite bursts from different foci at different moments may direct the immune response to antigens that are uniquely expressed or exposed in different tissue environments.

The rich set of biomarkers in this Atlas also provide essential information to study the dynamics of the more prevalent (shared) antibody specificities at an unprecedented depth and granularity. In chronic Chagas disease, antibody levels are maintained by the persistence of parasites and antigens (Zhang and Tarleton, 1999). Studying how the antibody repertoires fluctuate upon reduction of parasite loads or elimination, or during development of Chagas disease pathology is a clear path to discover markers for novel immunoassays.

The Human Chagas Antigen and Epitope Atlas is a reference resource that is freely accessible. The resource generated and described herein comprises the collection of antigenic regions of the *Trypanosoma cruzi* pangenome, as revealed by analyzing the anti-*T. cruzi* human antibody repertoires of 71 Chagas Disease patients. These individual antibody repertoires described in detail represent a foundational resource for the community that will serve as a major accelerator for the development of new diagnostics, serology-based immunoassays, vaccines, and to study the dynamics of adaptive immune responses at high resolution.

## ONLINE METHODS

### Array Designs

#### CHAGASTOPE-v1 Design used for antigen and epitope discovery

Two *T*.*cruzi* proteomes were used in this design: Sylvio X10 (*2*) and CL-Brener (*3*), both retrieved from TriTrypDB Release 5 (2016) (*4*). To produce a tiling display of peptides, we first merged sequences from both proteomes and parsed all proteins, removing those shorter than 16 amino acids and duplicates (based on protein ID). This resulted in 30,500 proteins: 10,832 for Sylvio X10 and 19,668 for CL-Brener. Next, we split proteins into peptides of length 16 with an offset of 4 amino acids between consecutive peptides (meaning that there was an overlap of 12 amino acids between those peptides). At this stage we removed duplicate peptide sequences thus each peptide was placed only once in the microarray. This set of 2,441,908 unique peptides is available as Supplementary Table S10 and as part of the submission to the ArrayExpress Database. Finally, we added other peptides from a number of sources to the CHAGASTOPE-v1 design, such as positive controls that corresponded to previously identified antigenic regions (*1*), and peptides from other pathogens that are seroprevalent in humans (e.g., cytomegalovirus). These positive controls were repeated 4 times across the array as peptides of length 15 with an offset of 1 amino acid to have a higher resolution for epitope mapping and to match the original conditions of past works. This, along with the trimming of a few peptides in the synthesis (see **Array Synthesis** below) resulted in a final array design containing 2,842,420 peptides, all of which were present only once in the microarray except for the positive controls. These peptides were assigned randomly to spots in the microarray.

#### CHAGASTOPE-v2 Design used for characterization of antigenic regions

Because the aim of this second design was to analyze a smaller subset of peptides in higher detail, we focused on the 9,547 antigenic regions found from the first design. We produced tiling displays of peptides spanning these regions, using peptides of length 16 with an offset of 1 amino acid (maximal resolution for epitope mapping), which resulted in 242,154 unique peptides. In this array design we also included additional peptide variants from the *T. cruzi* 231 strain (DTU TcIII) (*5*) that mapped to these regions as well as a number of detailed mutagenesis scans (AlaScan) of selected epitope (see main text). The final array design contained 392,299 addressable peptide spots. Peptides were assigned randomly to these spots. This design was used to drive synthesis of QX12 (12-plex) arrays where the same CHAGASTOPE-v2 design was replicated across all 12 sectors of the array (assayed individually).

### Human Serum Samples

Human serum samples from *T. cruzi*-infected patients and matched negative subjects used in this study were part of the collections of the Laboratorio de Enfermedad de Chagas, Hospital de Niños “Dr. Ricardo Gutierrez” (HNRG, Buenos Aires, Argentina) (AR; n=18); Fundación CEADES (Cochabamba, Bolivia) (BO; n=17); Protozoology Laboratory (LIM 49), Hospital das Clínicas, Faculdade de Medicina, Universidade de São Paulo (São Paulo, Brazil) (BR; n=18); Instituto Nacional de Salud Pública (Tapachula, Mexico) (MX; n=18); Instituto de Cardiología (Bucaramanga, Colombia) (CO; n=15); University of South Carolina (South Carolina, USA) (US; n=18). Human serum samples from patients with American Tegumentary Leishmaniasis (ATL) and matched negative subjects were from the Instituto de Patología Experimental, Universidad Nacional de Salta (IPE, Salta, Argentina) (LE; n=12). A list of samples and their code identifiers as used in this work are provided in Supplementary Table S2. Chagas Disease patients were in the asymptomatic chronic stage of the disease without cardiac or gastrointestinal compromise (age range: 15 to 96 years old, median: 48). Serum samples were collected from clotted blood obtained by venipuncture and analyzed for *T. cruzi*-specific antibodies with the following commercially or in-house kits: AR: Chagatest ELISA lysate (Laboratorios Wiener, Argentina), Chagatest HAI (Laboratorios Wiener, Argentina), Chagatest ELISA recombinant v4.0 (Wiener Lab, Argentina); BO: recombinant and lysate ELISA; BR: conventional in-house ELISA (confirmed by TESA Blot (*6*)); CO: ELISA (Chagatek, Organon, Argentina) and HAI (Chagatest, Weiner, Chile); MX: Bio-Rad Chagascreen Plus v4 (recombinant), Test ELISA para Chagas III (Grupo Bios, Chile), Accutrak Chagas Microelisa Test System (Laboratorio Lemos, Argentina), Accutrack Chagatek ELISA recombinante (Laboratorio Lemos, Argentina); US: Chagas Stat-Pak (Chembio, Medford, NY), Hemagen Chagas’ Kit ELISA (Hemagen Diagnostics Inc., Columbia, MD), and an in-house TESA-Blot (*6*). ATL samples were classified using an in-house ELISA based on crude antigen extracts from promastigotes and amastigotes of 3 different endemic *Leishmania* species and 2 reference strains (*7*). All procedures followed the Declaration of Helsinki Principles. Written informed consent was obtained from all individuals (or from their legal representatives), and all samples were decoded and de-identified before they were provided for research purposes. The procedures were approved by the following ethics committees: Hospital de Niños “Ricardo Gutierrez” (#CEI 14.14); Fundación CEADES (CE-CEADES-4-12-2018; IRB 0990-0279; FWA: 00024189) ; Comité de Etica en Investigación, Instituto Nacional de Salud Pública, Mexico (CI: 1369, Registro ante CONBIOÉTICA: 17CEI00420160708, Registro ante COFEPRIS: 13 CEI 17 007 36, FWA: 00015605), Comité de Ética en Investigación FOSCAL, Bucaramanga, Colombia (CEI 21/11/14) and Fundación Cardioinfantil (Acta 512/2015), the study sites of the CHICAMOCHA3-equity trial; Baylor College of Medicine (Houston, TX, USA) (#H-35471 and #H-32321), the Gulf Coast Regional Blood Center (Houston) (#13-002), and the South Texas Tissue and Blood Center (San Antonio, TX, USA); and Comisión Provincial de Investigaciones Biomédicas, Ministerio de Salud Pública, Gobierno de la Provincia de Salta, Argentina (Expte 321 -136934/2018). Samples from LIM 49 were part of an older collection of samples (8-10 years) and qualify as secondary research use of biospecimens for which informed consent was not required. These fall into exemption #4 in the list of exemptions for the requirement of informed consent developed by the Office for Human Research Protections (OHRP), US Department of Health & Human Services, as it did not involve new recruitment of human participants and samples did not include any direct identifier.

### Array Synthesis

The CHAGASTOPE array designs were synthesized at Roche Sequencing Solutions (Peptide Lab, Madison WI, now Nimble Therapeutics) with a Roche Sequencing Solutions Maskless Array Synthesizer (MAS) by light-directed solid-phase peptide synthesis using an amino-functionalized support (Greiner Bio-One) coupled with a 6-aminohexanoic acid linker and amino acid derivatives carrying a photosensitive 2-(2-nitrophenyl) propyloxycarbonyl (NPPOC) protection group (Orgentis Chemicals). Amino acids (final concentration 20 mM) were pre-mixed for 10 min in N,N-Dimethylformamide (DMF, Sigma Aldrich) with N,N,N’,N’-Tetramethyl-O-(1H-benzotriazol-1-yl)uronium-hexafluorophosphate (HBTU, Protein Technologies, Inc.; final concentration 20 mM) as an activator, 6-Chloro-1-hydroxybenzotriazole (6-Cl-HOBt, Protein Technologies, Inc.; final concentration 20 mM) to suppress racemization, and N,N-Diisopropylethylamine (DIPEA, Sigma Aldrich; final concentration 31mM) as base. Activated amino acids were then coupled to the array surface for 3 min. Following each coupling step, the microarray was washed with N-methyl-2-pyrrolidone (NMP, VWR International), and site-specific cleavage of the NPPOC protection group was accomplished by irradiation of an image created by a Digital Micro-Mirror Device (Texas Instruments), projecting 365 nm wavelength light. Coupling cycles were repeated to synthesize the full in silico-generated peptide library. Coupling cycles were limited to avoid extremely long synthesis times, which had the consequence of trimming some peptides in our design by a few amino acids (usually peptides where a single amino acid appeared many times). This occurred in 0.5% of the peptides in the first design and 1.4% of the peptides in the second one, with an average of 1.5 and 1.7 amino acids trimmed in each case respectively. Because this was a rare event, because the trimming removed only one or two amino acids, and because we also smoothed the signal data using a rolling median technique (see below), we assumed this trimming had no substantial impact on analysis of the data.

### Array Assays

For the antigen and epitope discovery screening using the CHAGASTOPE-v1 design we produced and assayed 28 1-plex high-density peptide arrays. For the epitope characterization, mapping and seroprevalence study using the CHAGASTOPE-v2 design, we produced and assayed 12 (twelve) 12-plex sectorized high-density peptide arrays. Supplementary Table S1 provides an overview of 1-plex and 12-plex arrays used in this work, while Supplementary Table S3 provides a list of arrays slides and samples used in each assay.

### Sample Binding and Detection

Prior to sample binding, final removal of side-chain protecting groups was performed in 95% trifluoroacetic acid (TFA, Sigma Aldrich), 0.5% Triisopropylsilane (TIPS, TCI Chemicals) for 30 min. Arrays were incubated twice in methanol for 30 s and rinsed four times with reagent-grade water (Ricca Chemical Co.). Arrays were washed for 1 min in TBST (1× TBS, 0.05% Tween-20), washed twice for 1 min in TBS, and exposed to a final wash for 30 s in reagent-grade water.

Serum samples were diluted 1:100 in binding buffer (0.01M Tris-Cl, pH 7.4, 1% alkali-soluble casein, 0.05% Tween-20) and bound to arrays overnight at 4°C. After sample binding, the arrays were washed three times in wash buffer (1× TBS, 0.05% Tween-20), 10 min per wash. Primary sample binding was detected via Alexa Fluor® 647-conjugated goat anti-human IgG secondary antibody (Jackson ImmunoResearch #109-605-098). The secondary antibody was diluted 1:10,000 (final concentration 0.1 ng/μl) in the secondary binding buffer (1x TBS, 1% alkali-soluble casein, 0.05% Tween-20). Arrays were incubated with secondary antibody for 3 h at room temperature, then washed three times in wash buffer (10 min per wash), washed for 30 sec in reagent-grade water, and then dried by spinning in a microcentrifuge equipped with an array holder.

Fluorescent signal of the secondary antibody was detected by scanning at 635 nm at 2 μm resolution and 15% gain, using an MS200 microarray scanner (Roche NimbleGen). Scanned array images were analyzed with proprietary Roche software to extract fluorescence intensity values for each peptide.

### Data Analysis. Normalization, quality control and removal of outliers (smoothing)

#### CHAGASTOPE-v1 - Quality control

All experiments were performed in duplicate (same biological sample, duplicate assays on independent array slides). We performed quality control of each pair of replicates for each sample using Bland-Altman (MA) plots and reciprocal signal plots. All replicate array assays showed excellent overall reproducibility (see Supplementary Figure S2). As another step of quality control, we analyzed the replicas of the positive controls we placed in the array (these were sections of known antigenic proteins that had their peptides repeated 4 times in the microarray design).

For this we used their normalized signals (see below) and we observed excellent overall reproducibility between the replicas, both intra- and inter-array (see Supplementary File S1).

#### CHAGASTOPE-v1 - Quantile normalization

To compare data across experiments, we normalized array data using quantile normalization. Because this method requires similar statistical properties of the underlying distributions, we performed two sets of quantile normalizations, one for the assays using Chagas-positive samples, and one for the assays using negative samples (including those from Leishmania positive serums, swhich produced signal ditributions similar to the Chagas-negative samples). We treated replicas as independent samples, resulting in a normalization across 12 array sets for the Chagas-positive samples and a normalization across 16 array sets for the rest. Normalization was performed in R using *normalize*.*quantiles* from the package *preprocessCore*.

#### CHAGASTOPE-v1 - Smoothing and replicas

To remove outliers, we used two methods combined: a rolling median smoothing procedure and an average via replicas. First, we assigned the normalized signal of each peptide to the corresponding parent protein sequence(s). This was done once per serum sample per replica. Next, we used the *rollmedian* function in the R package *zoo* to calculate the rolling median along the protein sequence. We used a window size of 3, meaning that the smoothed signal for each peptide was the median of itself and its two neighboring peptides in the same protein/replica/serum (for peptides at the edges of the protein sequences we added a 0 as the signal of the non-existing neighboring peptide). Next, we combined data from the two replicas to calculate their average and standard deviation, resulting in the final data set that was analyzed and described herein. In this final data set each peptide had 14 associated signal values (6 from Chagas-positive samples, 6 from Chagas-negative samples, 1 from a Leishmaniasis sample and 1 from a Leishmaniasis negative sample).

#### CHAGASTOPE-v2 - Quality control, quantile normalization and smoothing

In the CHAGASTOPE-v2 arrays we followed similar steps as in the first design. **Quality control:** we analyzed each pair of replicates for each sample using Bland-Altman (MA) plots and reciprocal signal plots (see Supplementary Figure S3). **Quantile normalization:** In these 12-plex assays, one microarray slide contained 12 sectors, which were assayed separately, hence for all data-analysis purposes 1 array sector was treated as one 1 array data set. Quantile normalization was thus performed for 142 assays (2 replicas for each of 71 individual serum samples). **Smoothing and replicas:** The smoothing and combining of replicas was done exactly as in CHAGASTOPE-v1.

### Data Analysis. Definition of antigenic peaks and regions

#### CHAGASTOPE-v1 - Antigenicity threshold

To define this threshold, we analyzed the normalized signals (before smoothing) for peptides in the 12 Chagas-positive pooled samples (excluding the Leishmania positive and negative pools) and calculated their mode and standard deviation. We then looked at the protein profiles (the smoothed signals) and analyzed the dispersion of the “noise”, meaning how high were the signals for the healthy pools. Since this was a discovery screening, we wanted to be very conservative in this choice, making sure to select regions that were truly antigenic. All this, coupled to the amount of space available in our second design, led to us using an antigenicity threshold of 10,784.80 arbitrary fluorescence units (mode plus 4 standard deviations).

#### CHAGASTOPE-v1 – Peaks

For the discovery screening, we defined as “peak” a group of two or more consecutive peptides with signals greater than the antigenicity threshold. Because we were interested in the discovery of *T. cruzi* antigens and epitopes, we also required each peak to have a maximum signal in a Chagas-positive sample that was at least five times higher than the corresponding maximum signal in the negative samples for those peptides.

#### CHAGASTOPE-v1 – Regions

Antigenic Regions result from merging of neighboring peaks. For each peak we noted the position in its protein of the first and last peptides. We then expanded this range by moving the start of the peak 16 amino acids to the left and the end of the peak 16 to the right to ensure capturing the whole peak. Then, if two or more of these new “wide peaks” overlapped between each other they were all merged into one. This resulted in 9,547 antigenic regions across both proteomes.

#### CHAGASTOPE-v1 - Clusters of regions

In our analyzed proteomes there were identical proteins or different proteins sharing significant sequence similarity over a domain or defined sequence region. This was either because they belonged to the same protein family or because the protein was present in both CL-Brener (Esmeraldo and Non-Esmeraldo haplotypes) and Sylvio X10 proteomes. This similarity resulted in several antigenic regions with very similar or identical sequences, which can distort the conclusions drawn from the data. To reduce redundancy, we clustered antigenic regions by sequence similarity using *blastp* (BLAST 2.2.31+) (*8*). The all vs all comparison across all 9,547 regions was run with the following *blastp* command options and parameters:

~~~
-outfmt ‘6 qseqid sseqid pident length mismatch gapopen evalue bitscore qseq sseq sstart send’ -word_size 2 -comp_based_stats 0 -max_target_seqs 50000 -matrix BLOSUM80
~~~

We kept only matches with a percentage of identical amino acids (pident) of at least 80% and a match length of at least 75% of the length of the shortest region in the match.

Using these matches, we computed a distance matrix where distance was 1 - (pident/100) and then applied a single-linkage hierarchical clustering method. The resulting tree was cut at a cutoff of 0.2 (1 - pidentThreshold), resulting in 3,868 Distinct Antigenic Regions.

#### CHAGASTOPE-v2 - Antigenicity threshold

To determine the antigenicity threshold in this experiment, we set to recreate the results obtained in the discovery screening using virtual sample pools. The signal of a virtual pool for a given peptide was the highest signal for that peptide amongst the individual serums that were part of that pool (and were now being analyzed individually). We then compared the antigenicity of these virtual pools against the antigenicity from our original pools in all clusters of antigenic regions using ROC curves. The goal was to predict the antigenicity in the original pools using the information from the corresponding virtual pools. An original pool was antigenic if it surpassed the 10,784.80 threshold, but for the virtual pools we analyzed possible thresholds of the formula mode plus X standard deviations, where X ranged from 1 to 4 in steps of 0.1 (using the mode and standard deviation from the second design). In the end the best threshold was 5,814.81 (mode plus 2.4 standard deviations) with an AUC of 0.83.

#### CHAGASTOPE-v2 - Comparison between strains

We analyzed the 33 samples from individuals that were used at the CHAGASTOPE-v1 discovery stage as pooled samples. We focused initially on these samples because they were used both to select reactive peptides for the CHAGASTOPE-v2 design and in the individual serological profilings of CHAGASTOPE-v2. For each individual sample, we assessed the number of reactive peptides that were exclusively present in the CL-Brener or Sylvio X10 proteomes. To define if a peptide belonged to CL-Brener or Sylvio (or to both), we looked at each of the peptides from the antigenic regions present in CHAGASTOPE-v2 and mapped these to their cognate proteins. Peptides that were mapped to antigenic regions in both CL-Brener and Sylvio X10 were excluded from this analysis. Next, we counted the number of non-redundant reactive peptides that were exclusive for each strain and for each of the individual serum samples. When calculating the signal for the non-redundant peptides we kept the highest signal between the peptides with the same sequence. Because the CL-Brener proteome is larger (almost double the size of the Sylvio X10 proteome) we standardized the number of reactive non-redundant peptides using two sets of z-scores, one for each strain.

We also carried out a similar analysis using all 71 individual samples from CHAGASTOPE-v2. This is presented in Supplementary Figure S5. In this case, however, interpretation of the relative reactivity of these samples against the two *T. cruzi* strains has to be done with care, because 38 of these samples were not used on proteome-wide v1 arrays, and hence were not used to select peptides for inclusion in the CHAGASTOPE-v2 arrays.

#### CHAGASTOPE-v2 - Single residue mutational scannings

From the peaks found in the CHAGASTOPE-v1 primary screen we selected those with antigenicity in at least two pools and a maximum signal of at least 21,500 fluorescence units (these parameters were a consequence of the space assigned in our second design for this experiment).

This resulted in 1,445 antigenic peaks from 977 proteins. For each of these peaks we selected its best peptide and added its 4 neighboring amino acids (2 to each side). This resulted in a set of sequences of 20 amino acids, from which 789 were distinct sequences.

In this study we analyzed all these sequences, although some different peaks are actually very similar with only residue variants. Therefore, and to clarify presentation, peaks were clustered by similarity into 232 significantly different sequences. To accomplish this, we first calculated similarity across sequences using BLOSUM 80 and the function parSeqSim from R package “protr” (*9*). Then, we grouped the similar sequences in clusters, trying all possible numbers of clusters and selecting the optimal number according to a silhouette analysis scanning the performance. This resulted in 144 clusters which were later manually curated, where some clusters were splitted while others were merged, resulting in the final 232 significant sequences. For each cluster we only present the result for the most antigenic peak sequence.

Mutational scanning of candidate epitopes was performed on a set of 5 overlapping peptides (16mers) which in concert cover a sequence of 20 aminoacid residues. This is explained visually in Figure 7 and in Supplementary Figure S6. In a glance, the best 16mer (max signal) in a selected antigenic peak was labeled as the central peptide (position 0), then the 4 additional neighboring peptides were labeled as their position relative to the central peptide, meaning positions -2, -1, +1, and +2. This resulted in 2,914 unique peptides. The antibody-binding signal of each of these 5 peptides correspond to the non-mutated (wild-type) versions of an epitope. To analyze the contribution of each residue in this sequence to the binding of specific antibodies, we generated all possible alanine variants (*ie*. the replacement of each original amino acid in the sequence for alanine) from those 5 original peptides. In case the original amino acid was an alanine, glycine was used as replacement. Since the peptides were 16mers (16 residues of length) we derived in 80 mutated sequences for each peak, which added to the 5 original peptides resulted in a total of 85 peptides for each peak. A total of 45,519 unique peptides were generated for this mutational analysis.

Supplementary Figure S6 shows a schematic view of this procedure for one antigen. Each residue from each peak sequence was analyzed individually. The difference with the mean signal of the peptides that contained the non-mutated (or original) residue and those that contained the residue replaced by alanine or glycine, as appropriate, was calculated for each serum or several. This difference in each residue defines a profile change. See Supplementary File S6 for the full results of the example of Supplementary Figure S6. An arbitrary cut-off point was established at 1,300 to highlight residues with the greatest absolute change. Following this, sera were grouped according to the signal change profile of core residues (mean change > 1,300). Finally, a sequence-logo was constructed using a modified version of the R package “ggseqlogo” (*10*) where the height of the amino acid is directly proportional to the signal change for proper visualization.

## ABBREVIATIONS

DTU: Discrete Typing Unit

## ACKNOWLEDGEMENTS

The authors would like to acknowledge Santiago J Carmona (Swiss Bioinformatics Institute, University of Lausanne) for critical reading of the manuscript, and Centro Regional de Chagas, Hospital Independencia, Santiago del Estero, Argentina, and Renato Uncos for technical support.

## DATA AND MATERIALS AVAILABILITY STATEMENT (DAS)

Peptide array designs have been deposited in the European Array Express Archive at the European Molecular Biology Laboratory -- European Bioinformatics Institute (www.ebi.ac.uk/arrayexpress) under accession numbers A-MTAB-692, A-MTAB-693 (Array Designs).

Assay Data for CHAGASTOPE-v1 (*T. cruzi* Whole Proteome Array), corresponding to these 14 sera pools AR_PO, AR_NE, BO_PO, BO_NE, BR_PO, BR_NE, CO_PO, CO_NE, MX_PO, MX_NE, US_PO, US_NE, LE_PO, LE_NE), have been deposited in the European Array Express Archive under accession number E-MTAB-11651, containing raw and processed assay data. See also Supplementary Table S3.

Assay Data for CHAGASTOPE-v2 (Seroprevalence and Epitope Characterization), corresponding to 71 individual serum samples, have been deposited in the European Array Express Archive under accession number E-MTAB-11655, containing raw and processed assay data. See also Supplementary Table S3.

Due to their size, some Additional Supplementary Materials are available at this Figshare Collection (DOI: 10.6084/m9.figshare.19991021. These Materials include the following: Tables S10, S11, S12 and S13; Files S1, S2, S3, S4, S5, S6, S7, and S8.

Data can also be explored interactively at the https://chagastope.org website.

## CODE AVAILABILITY STATEMENT

Custom software used for data analysis are available at this GitHub Repository: https://github.com/trypanosomatics/The-Chagas-Disease-Antigen-and-Epitope-Atlas.

## ETHICS STATEMENT

This study was conducted in accordance with the Declaration of Helsinki. Written informed consent was obtained from all donors, except where noted (Exemption 4). This study was approved by the Ethical Committee of the Instituto de Investigaciones Biotecnológicas, Universidad Nacional de San Martín. Research did not involve interaction with the serum donors nor their identification.

## AUTHOR CONTRIBUTIONS

### CRediT - Contributor Roles

Conceptualization: ADR LEB FA; Data curation: ADR LEB FA; Formal analysis: ADR LEB; Funding acquisition: JA FA; Investigation: ADR LEB FA; Methodology: ADR LEB FA; Project administration: FA; Resources: JMR FT NK MSN JCV JA JDM FA; Software: ADR LEB; Supervision: FA; Validation: ADR LEB FA; Visualization: ADR LEB FA; Writing – original draft : ADR LEB FA; Writing – review & editing: ADR LEB JMR MSN FA

## FIGURES AND TABLES

In the manuscript produced for review purposes, all Figures and Tables are included in the main text, placed close to their first mention/reference.

