## Supplementary Figures for "A *Trypanosoma cruzi* Antigen and Epitope Atlas: deep characterization of antibody specificities in Chagas Disease patients across the Americas"

**This PDF file includes:**

- Supplementary Figures S1 to S12

**Other supplementary materials include the following:**

- Supplementary Tables S1 to S9 (additional Excel file).
- Supplementary Tables S10 to S13 (available at Figshare), as stand-alone files (ZIP compressed TSV files).
- Supplementary Files S1 to S8 (available at Figshare), as stand-alone files (PDF or Zip compressed TSV files).

Due to their size, some Additional Supplementary Materials are available at this Figshare Collection (DOI: 10.6084/m9.figshare.19991021). These Materials include the following: Tables S10, S11, S12 and S13; Files S1, S2, S3, S4, S5, S6, S7, and S8.

### FIGURES

Because we needed to display complete proteomes in the form of tiling peptides covering all proteins, it was necessary to optimize the use of the peptide array capacity. In a previous study for 457 *T. cruzi* proteins (Carmona et al., 2015) we fine mapped epitopes using overlapped peptides with a sliding window of 1 amino acid residue (maximum resolution). Here we performed simulations on those data where we increased the offset between overlapping peptides while measuring the success at mapping the epitopes of known antigens. Performance at this task was measured by the Area Under the ROC Curve. Performance was high and stable for offsets 1-3 (AUCs 0.825 -- 0.96) and showed only a slight decrease with an offset of 4 (AUC 0.76 -- 0.93). Greater spacing between peptides (less overlap) affected performance down to an AUC of 0.66 -- 0.79 for an offset of 10 (see Supplementary Figure S1).

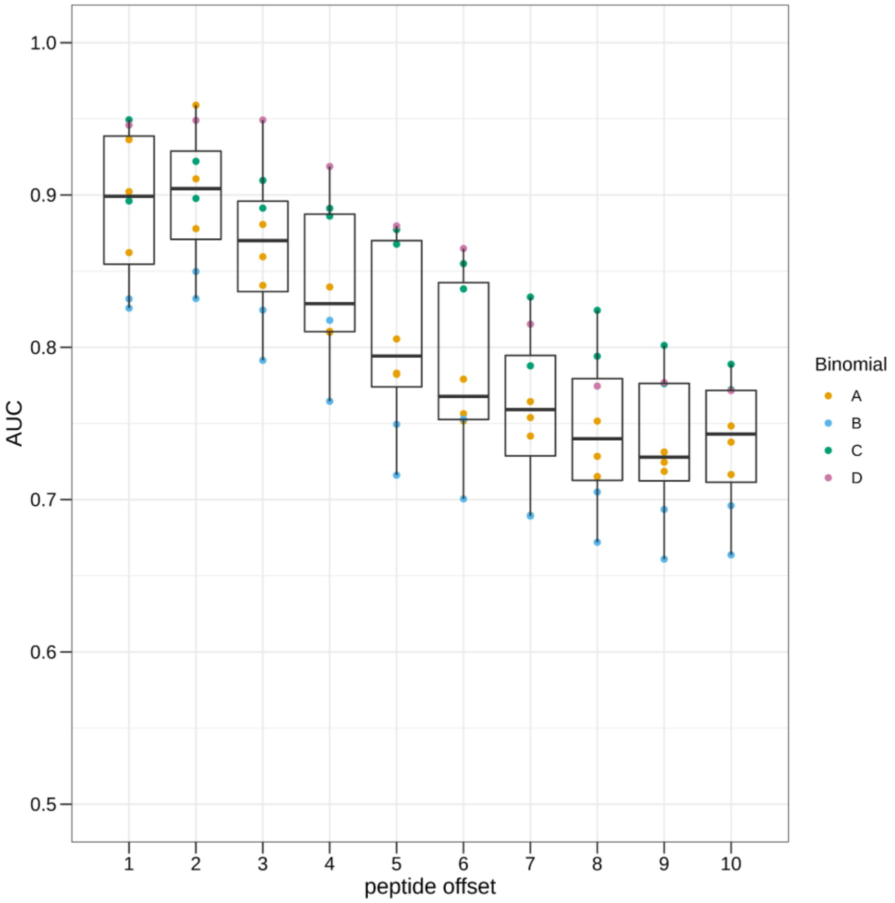


***Supplementary Figure S1. Performance at the task of mapping known linear epitopes using tiling peptide strategies.*** *Data from high-resolution peptide arrays (n=8) (Carmona et al 2015) were used to simulate different offsets. The original data contained maximal resolution epitope mapping assays (sliding window = 15 residues; offset = 1 residue). The simulated data was produced by skipping a different number of consecutive peptides in the sequence to produce lower resolution epitope mapping scenarios (skipping every other peptide produces offset = 2; skipping two consecutive peptides produces offset = 3, etc). In each simulation, we calculated the Area Under the ROC Curve (AUC) to measure the performance of a given offset at mapping the epitopes of known antigens.*

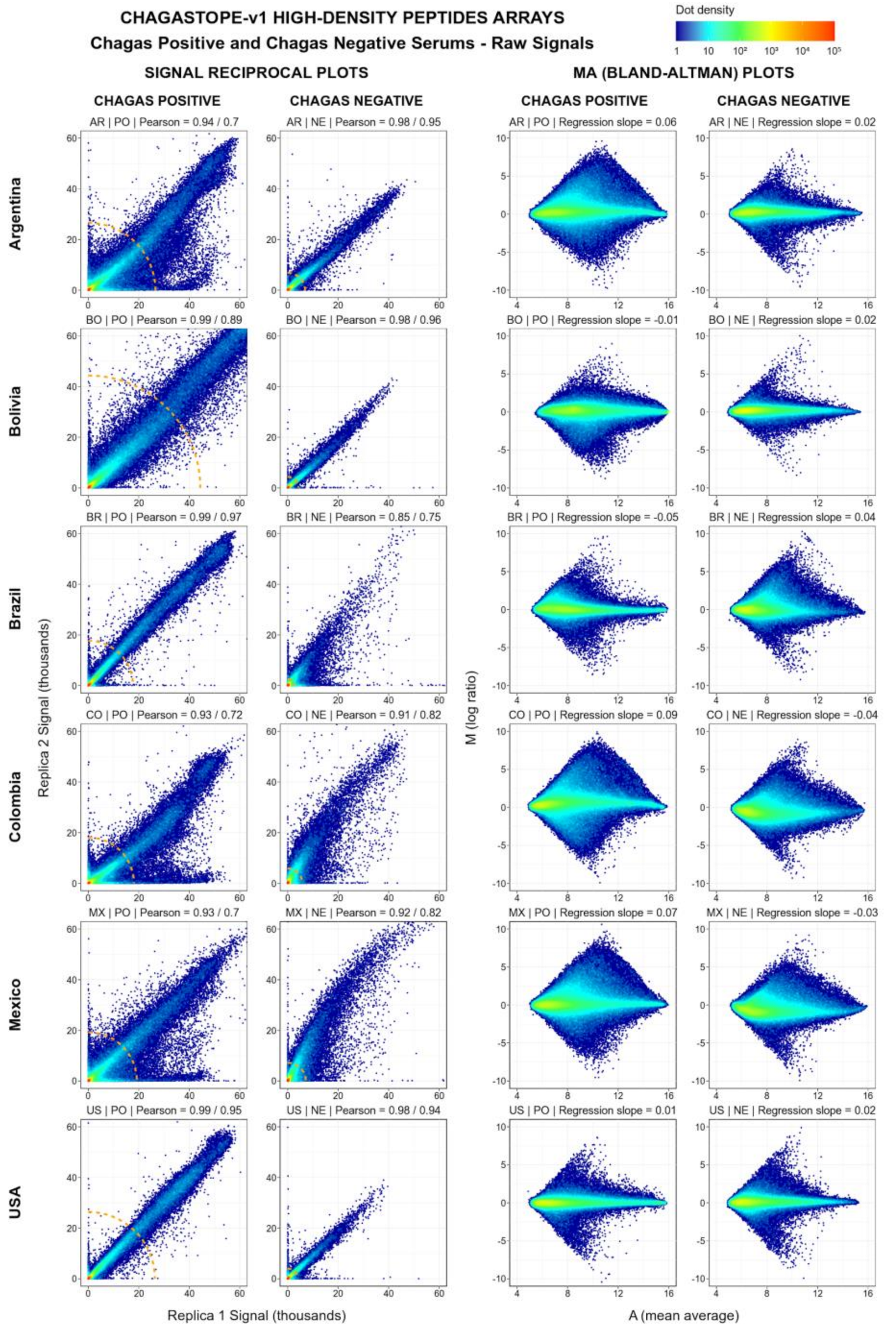


***Supplementary Figure S2a. Quality control of 1-plex microarray assays.*** *Each biological sample (serum pool) was assayed in duplicate (technical replicates). The signal correlation between technical replicates is shown both in reciprocal plots and in MA (Bland-Altman) plots. Each point represents one unique peptide or addressable array spot (n = 2,842,420). The density is shown in a color gradient (see key in figure). In the reciprocal plots, the raw signal data of one replicate is plotted against the second replicate and two Pearson scores are calculated: the first one using all peptides and the second one using only the ones with the top 1% signals, which are those outside the dashed orange line. In MA plots, each signal is replaced by its log2 and then the average signal of the two replicates (A) is plotted against the difference of the two signals (M).*

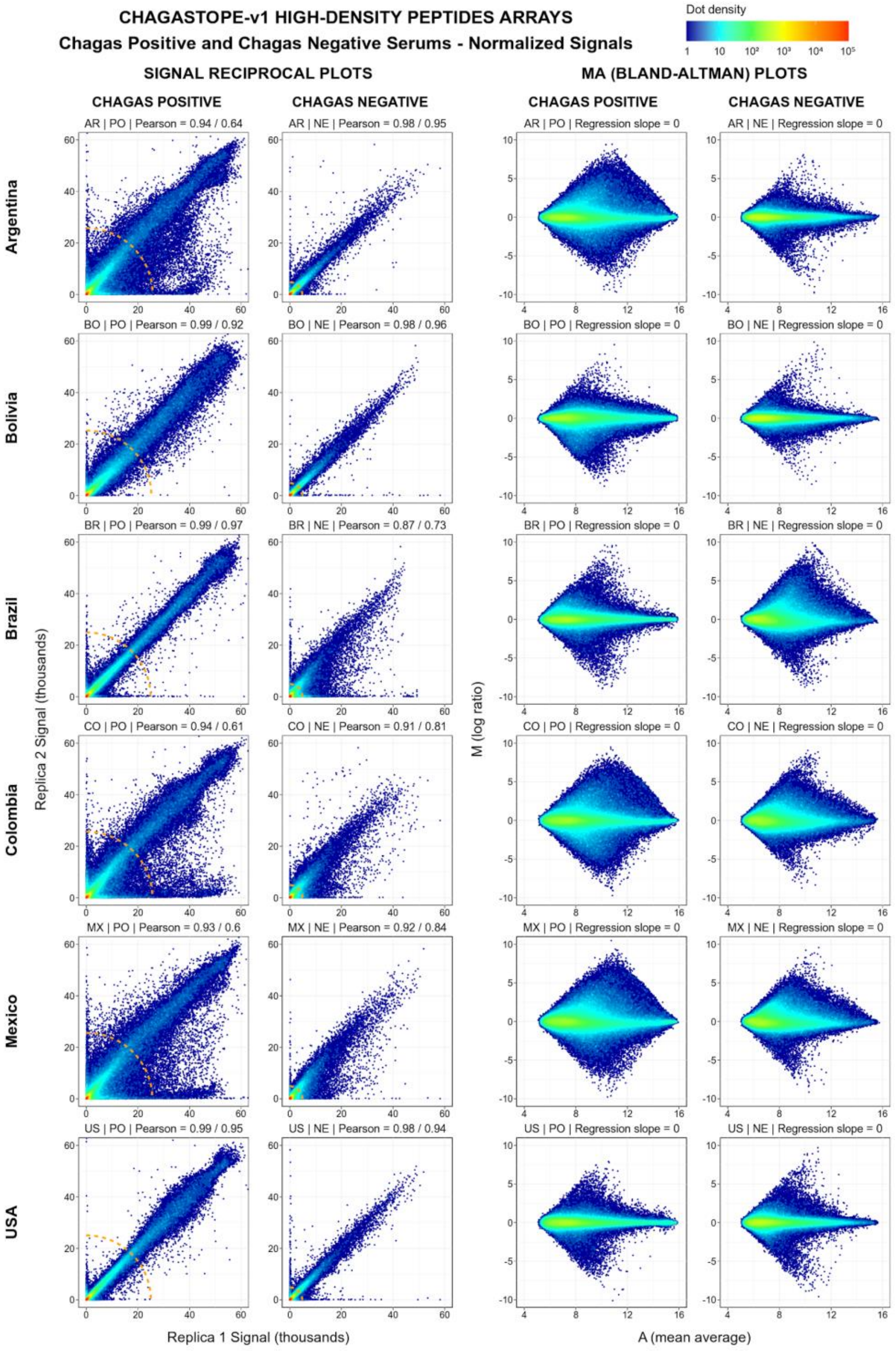


***Supplementary Figure S2b. Quality control of 1-plex microarray assays after normalization.*** *The legend is similar to that in Supplementary Figure S2a, but for the normalized signals.*

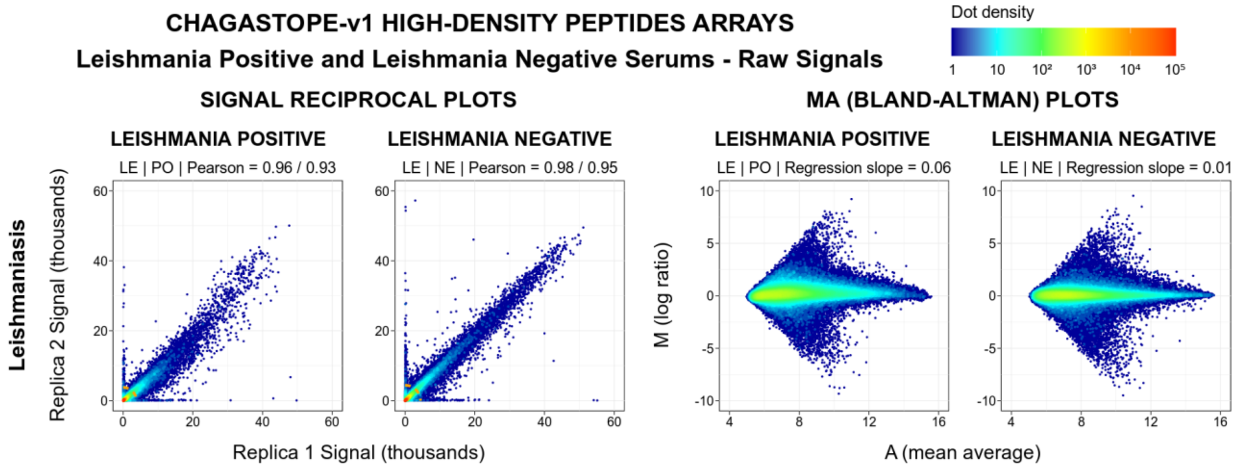


*__Supplementary Figure S2c. Quality control of 1-plex microarray assays for Leishmania.__ The legend is similar to that in Supplementary Figure S2a, but for the Leishmania positive and negative controls.*

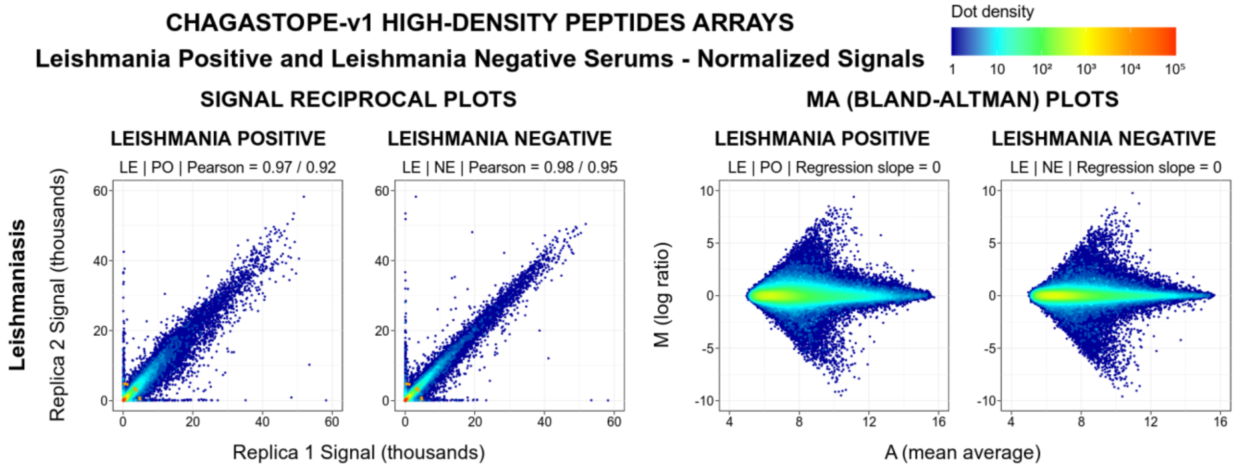


*__Supplementary Figure S2d. Quality control of 1-plex microarray assays for Leishmania after normalization.__ The legend is similar to that in Supplementary Figure S2c, but for the normalized signals.*

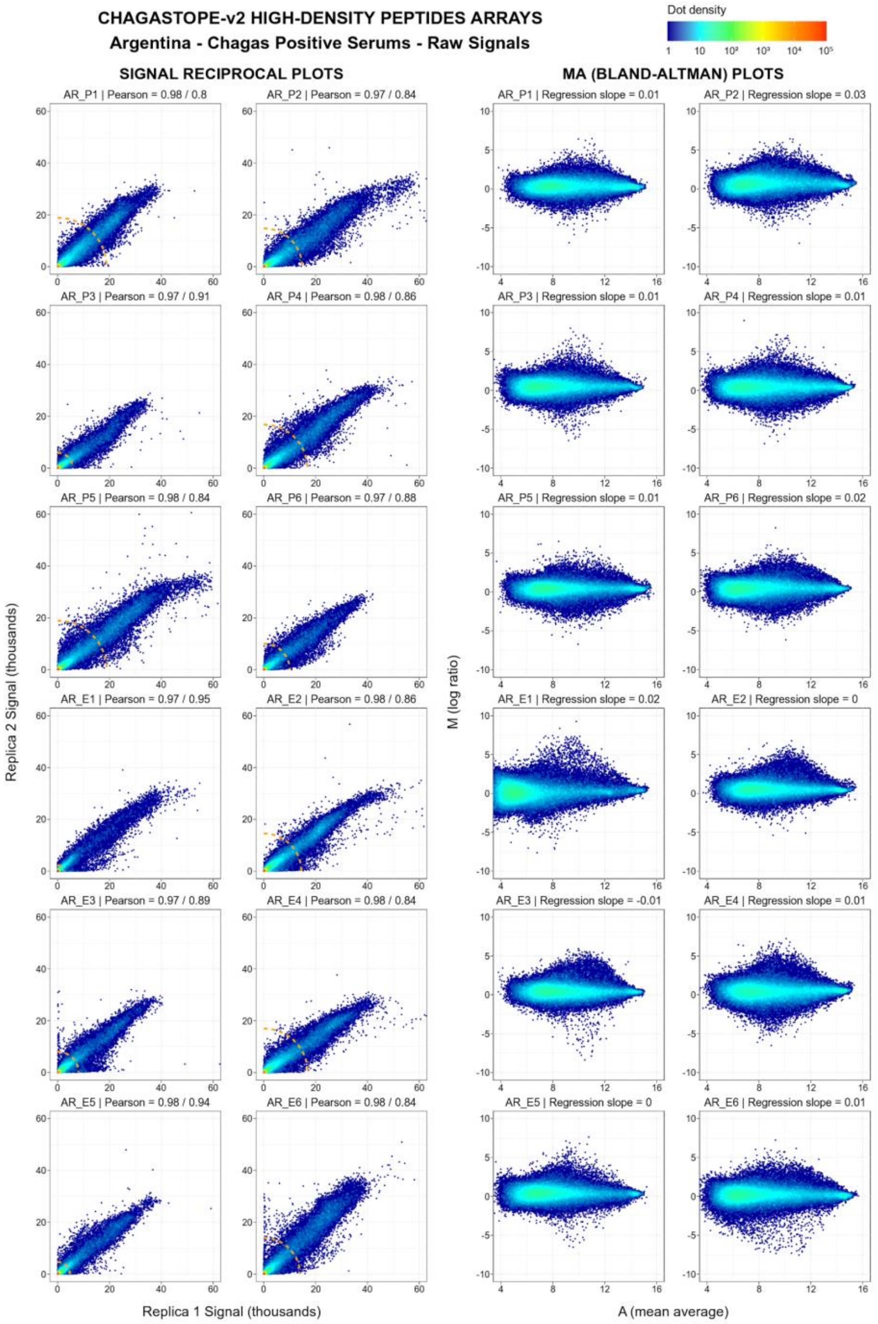


***Supplementary Figure S3a. Quality Control of 12-plex microarray assays for AR samples.*** *Each biological sample (serum sample from a single individual) was assayed in duplicate (technical replicates) in a separate 12-plex slide. The signal correlation between technical replicates is shown both in reciprocal plots and in MA (Bland-Altman) plots. Each point represents one unique peptide or addressable array spot (n = 392,299). The density is shown in a color gradient (see key in figure). In the reciprocal plots, the raw signal data of one replicate is plotted against the second replicate and two Pearson scores are calculated: the first one using all peptides and the second one using only the ones with the top 5% signals, which are those outside the dashed orange line. In MA plots, each signal is replaced by its log2 and then the average signal of the two replicates (A) is plotted against the difference of the two signals (M).*

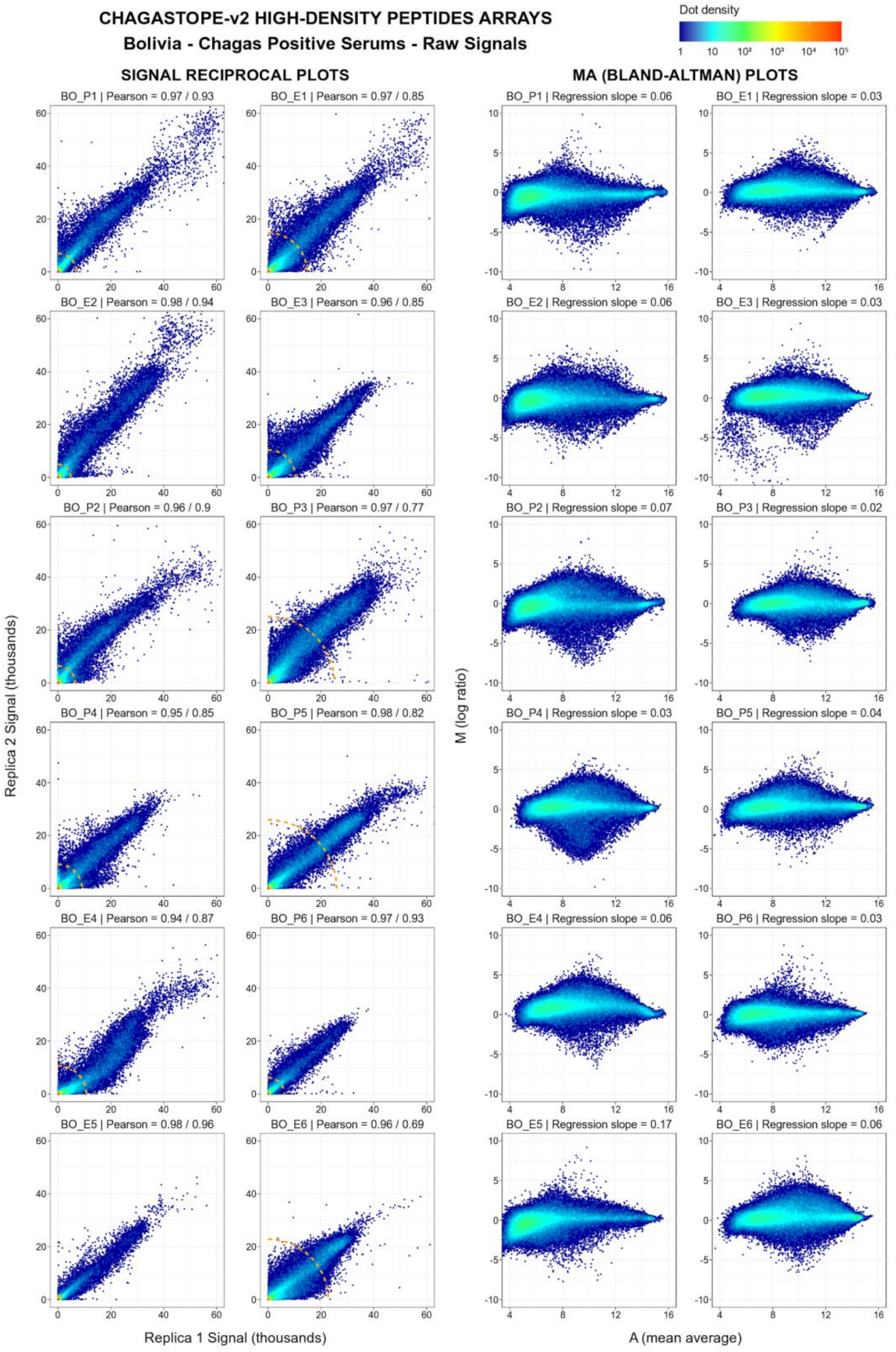


***Supplementary Figure S3b. Quality Control of 12-plex microarray assays for BO samples.***
*The legend is similar to that in Supplementary Figure S3a.*

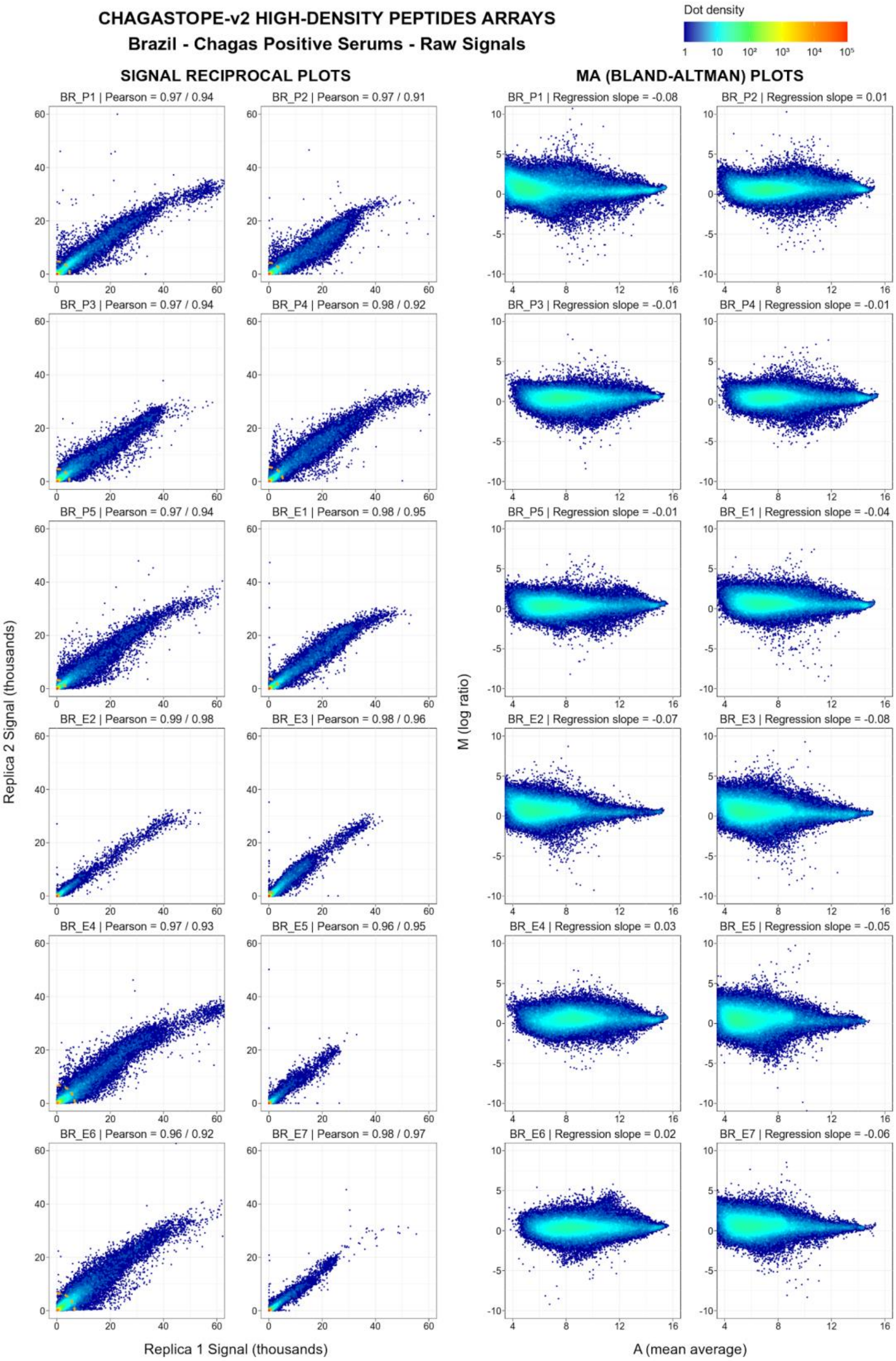


***Supplementary Figure S3c. Quality Control of 12-plex microarray assays for BR samples.***
*The legend is similar to that in Supplementary Figure S3a.*

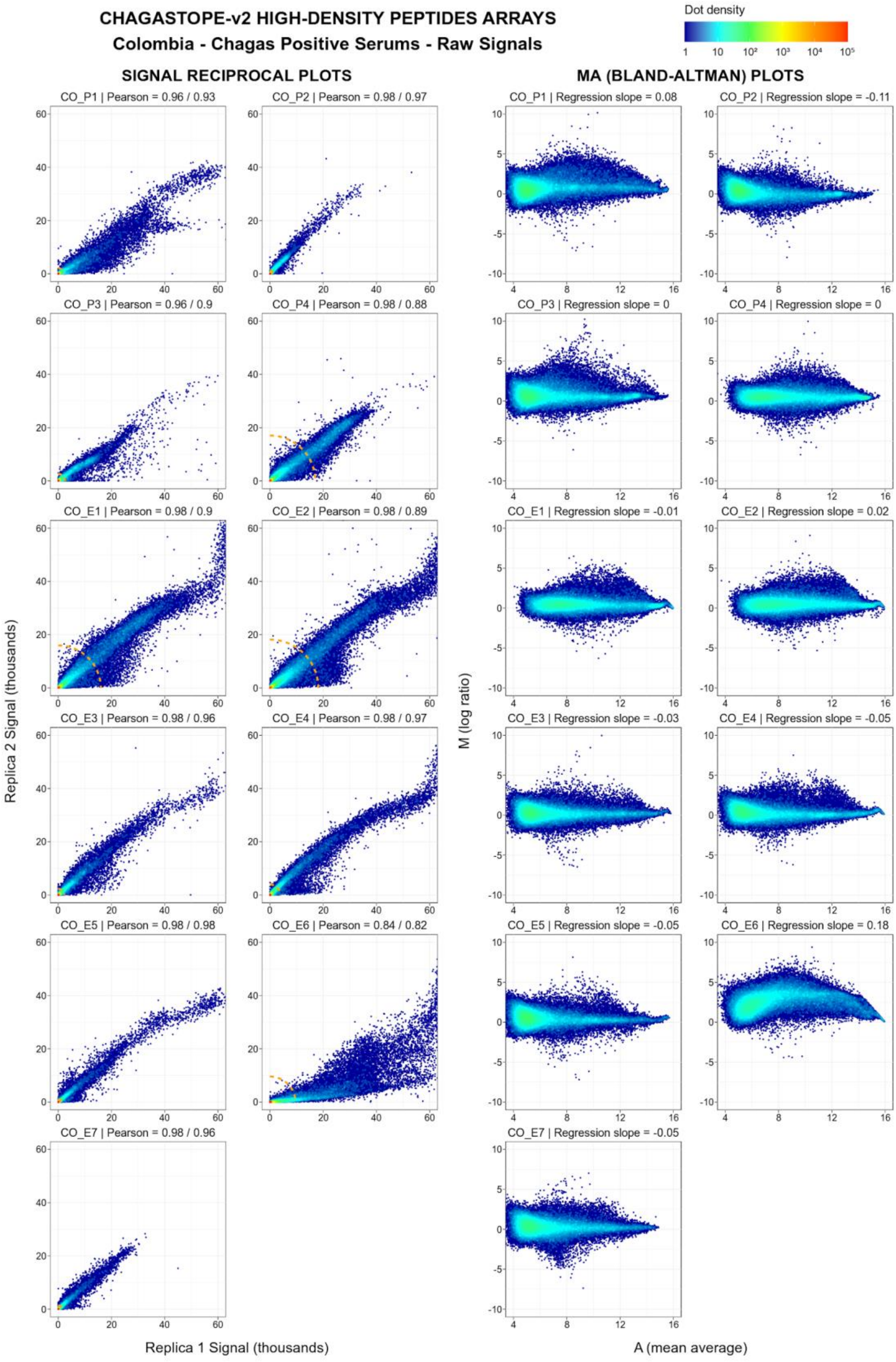


***Supplementary Figure S3d. Quality Control of 12-plex microarray assays for CO samples.***
*The legend is similar to that in Supplementary Figure S3a.*

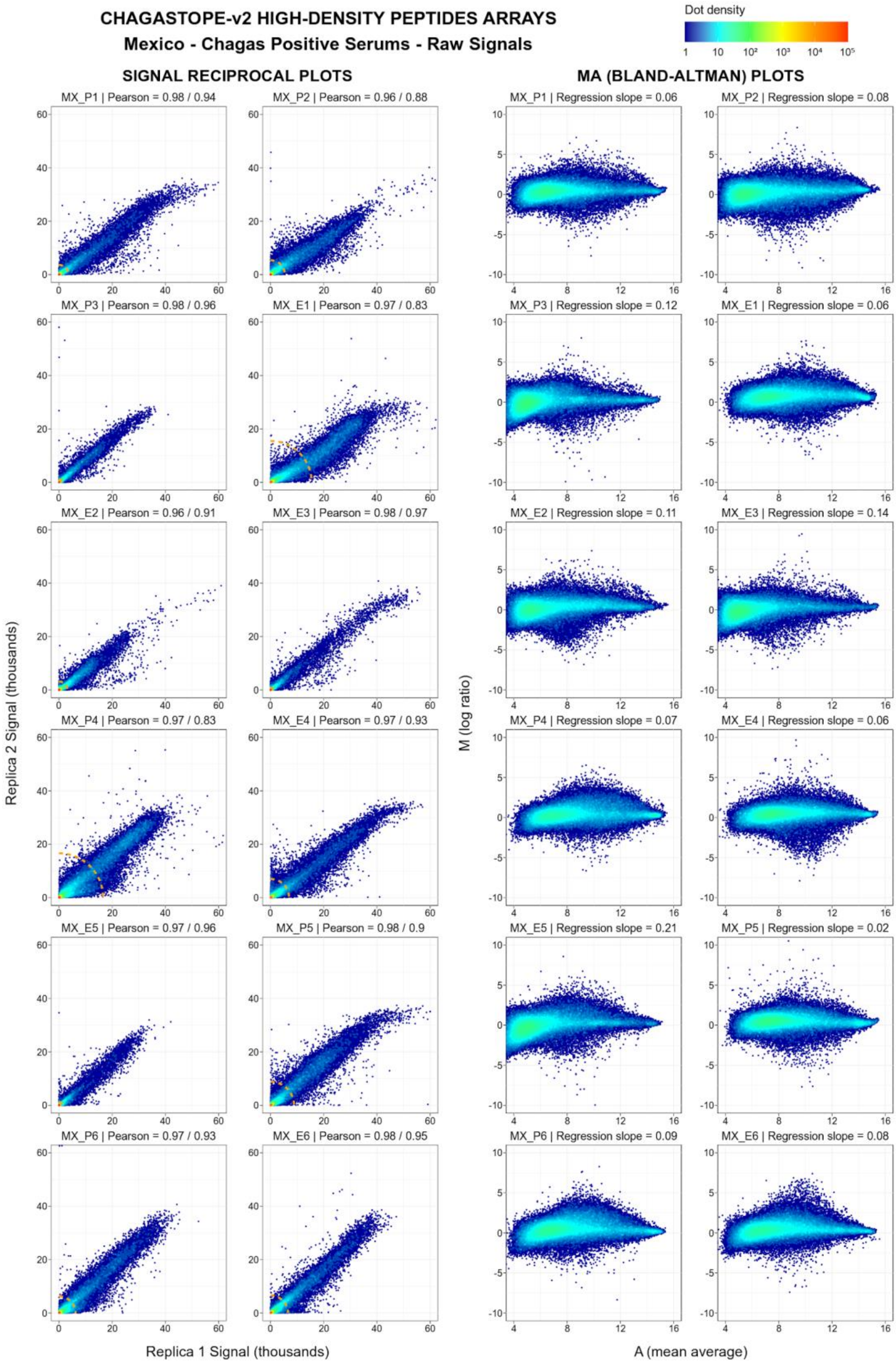


***Supplementary Figure S3e. Quality Control of 12-plex microarray assays for MX samples.***
*The legend is similar to that in Supplementary Figure S3a.*

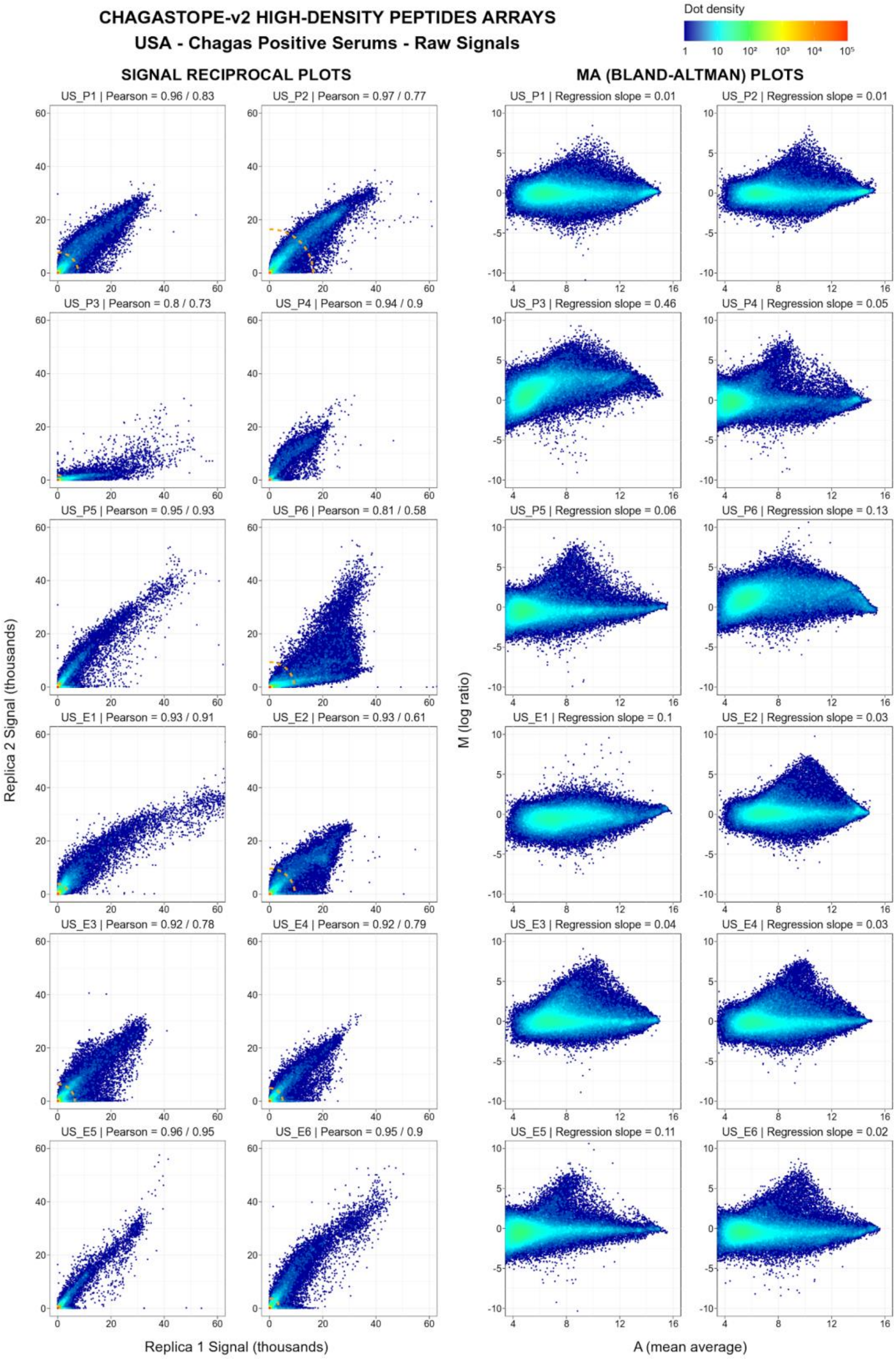


***Supplementary Figure S3f. Quality Control of 12-plex microarray assays for the US samples.*** *The legend is similar to that in Supplementary Figure S3a.*

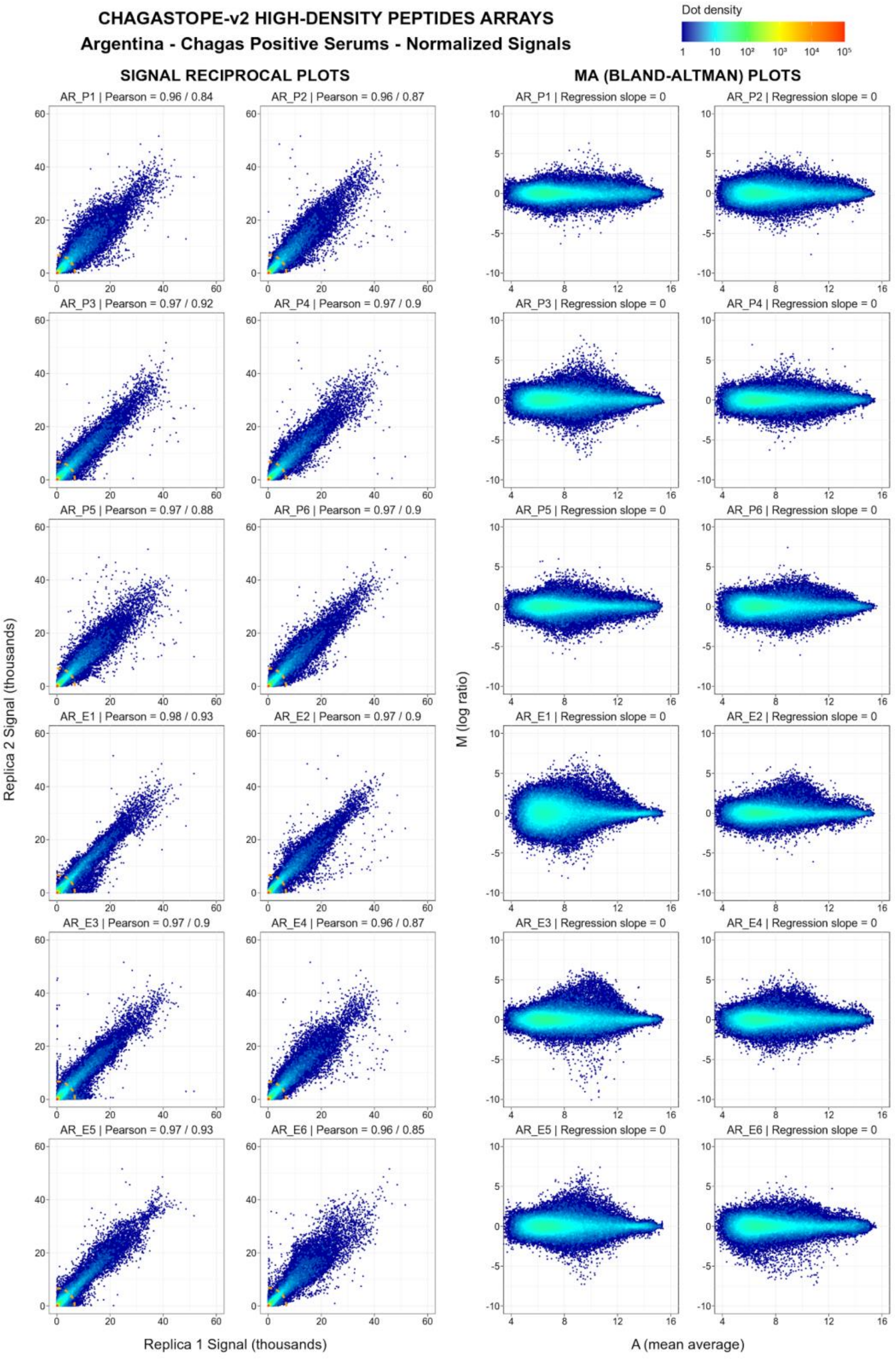


***Supplementary Figure S3g. Quality Control of 12-plex microarray assays for AR samples after normalization.*** *The legend is similar to that in Supplementary Figure S3a, but for the normalized signals.*

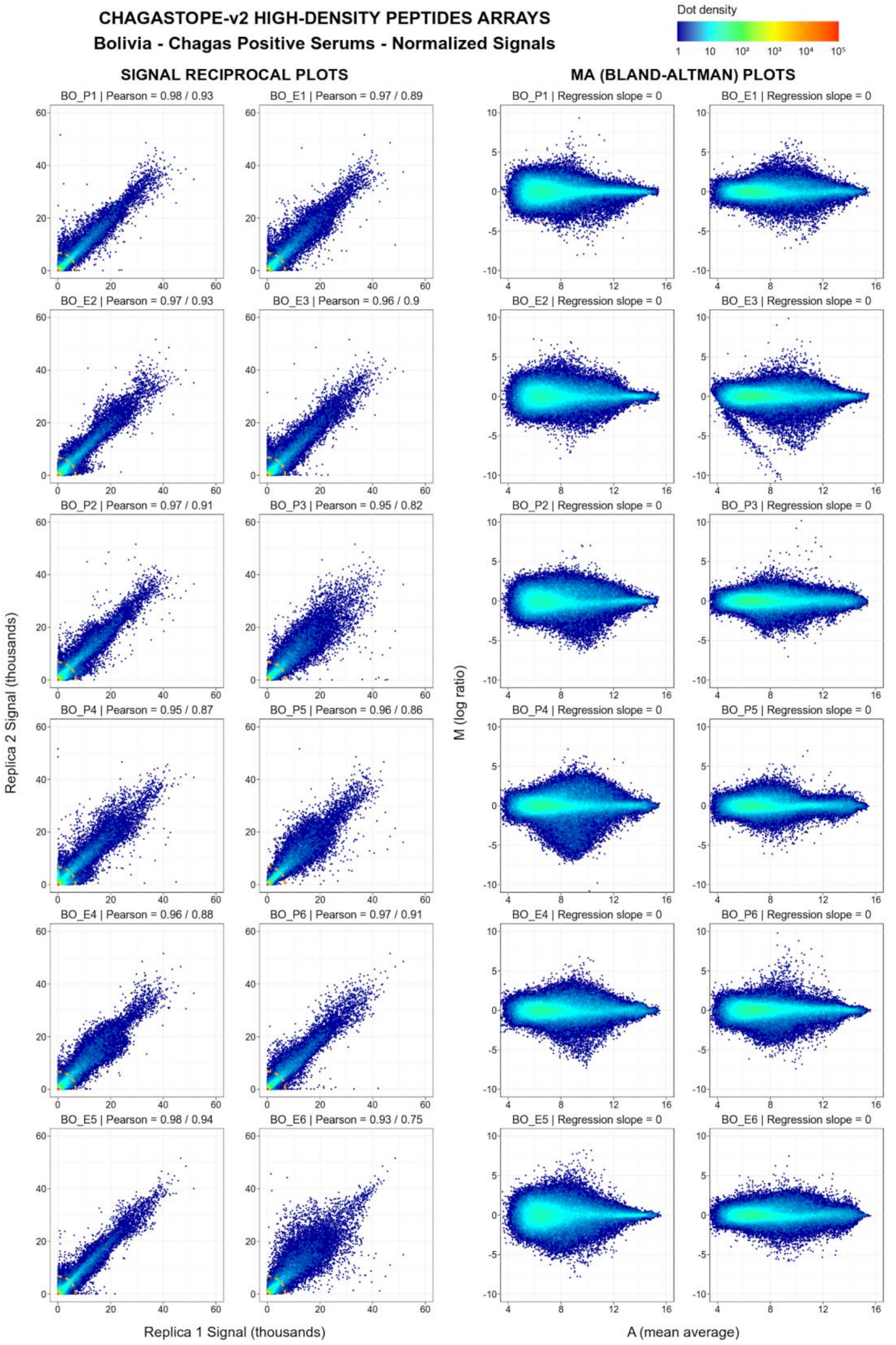


***Supplementary Figure S3h. Quality Control of 12-plex microarray assays for BO samples after normalization.*** *The legend is similar to that in Supplementary Figure S3a, but for the normalized signals.*

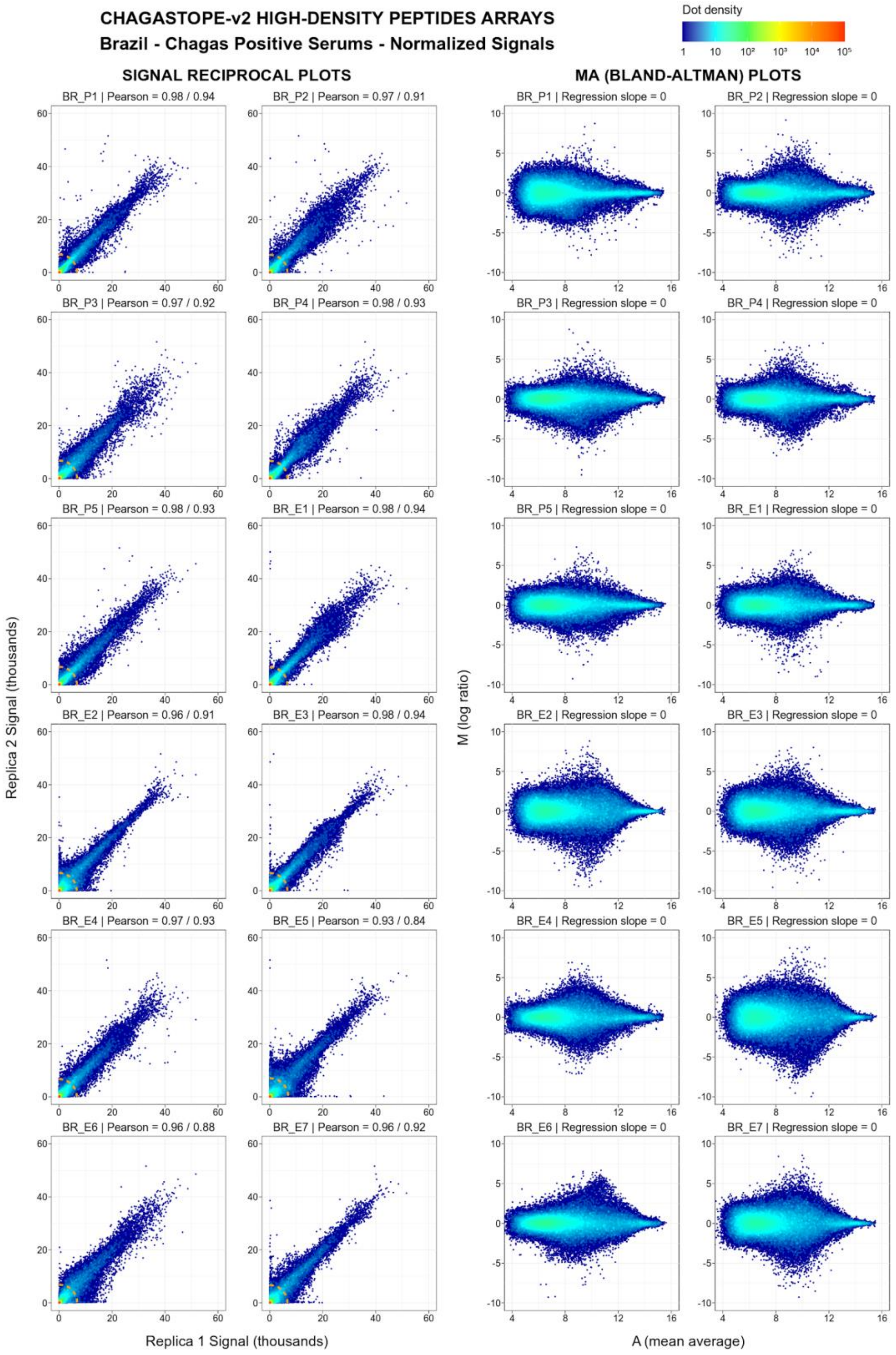


***Supplementary Figure S3i. Quality Control of 12-plex microarray assays for BR samples after normalization.*** *The legend is similar to that in Supplementary Figure S3a, but for the normalized signals.*

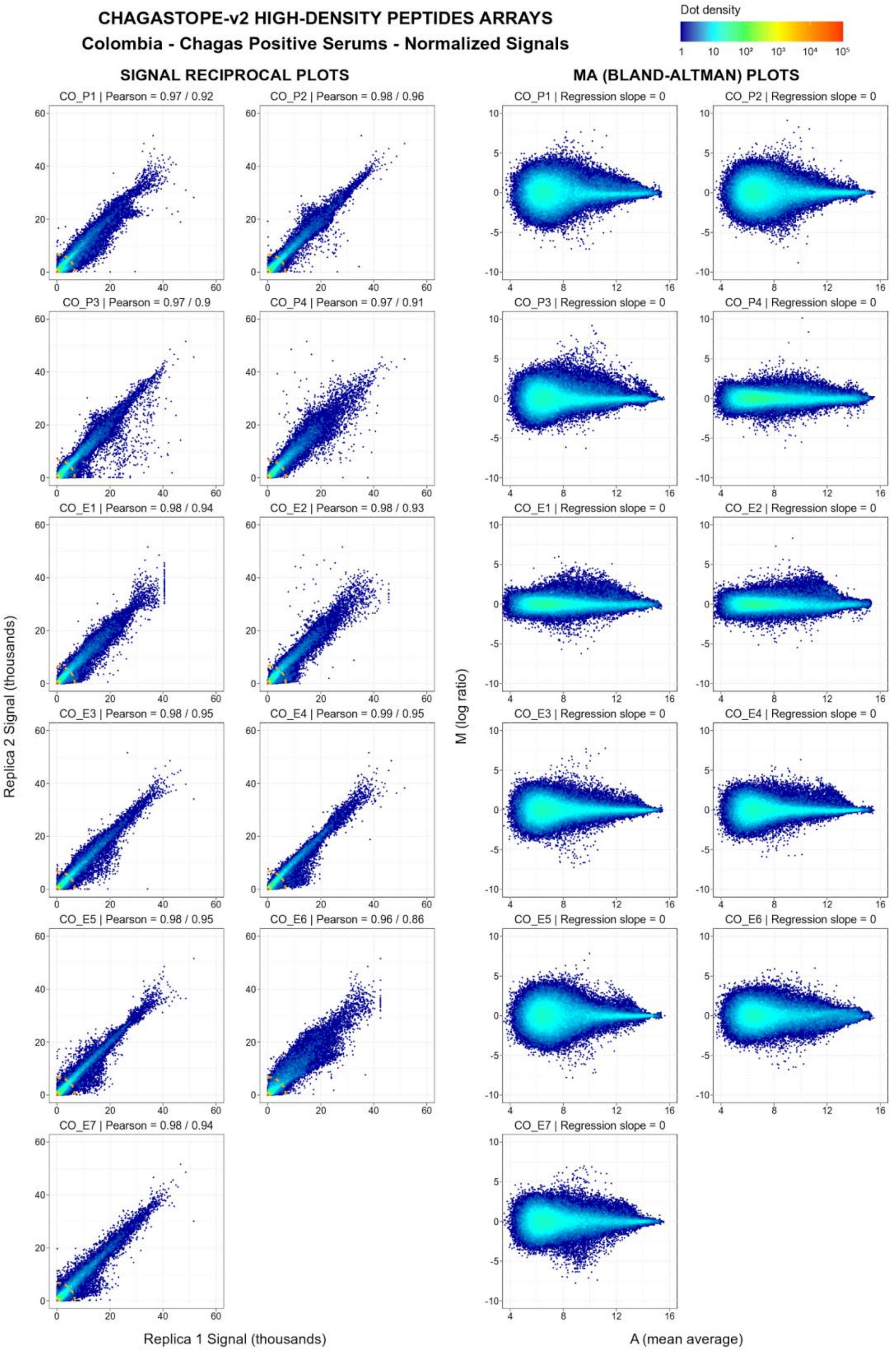


***Supplementary Figure S3j. Quality Control of 12-plex microarray assays for CO samples after normalization.*** *The legend is similar to that in Supplementary Figure S3a, but for the normalized signals.*

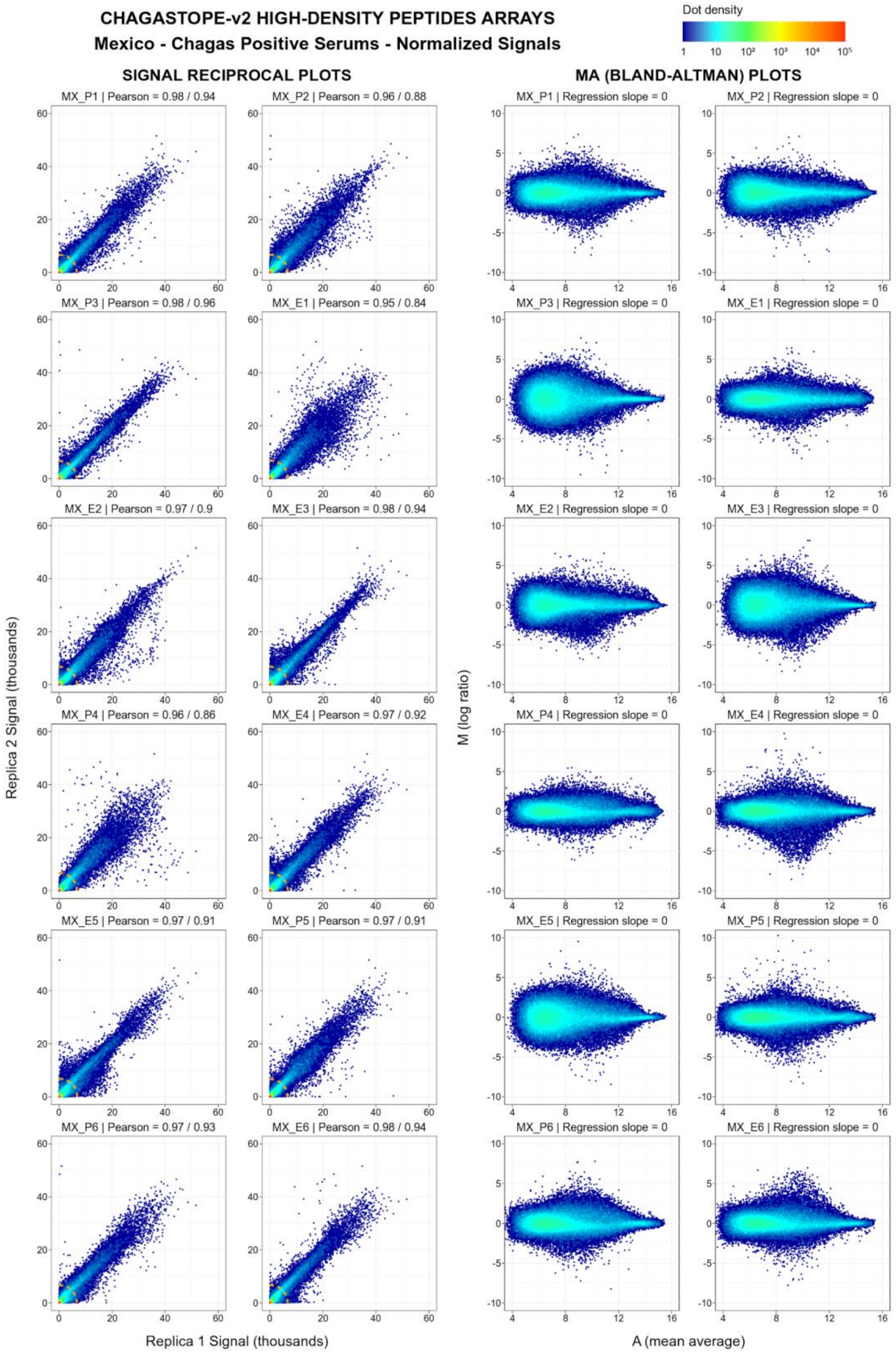


***Supplementary Figure S3k. Quality Control of 12-plex microarray assays for MX samples after normalization.*** *The legend is similar to that in Supplementary Figure S3a, but for the normalized signals.*

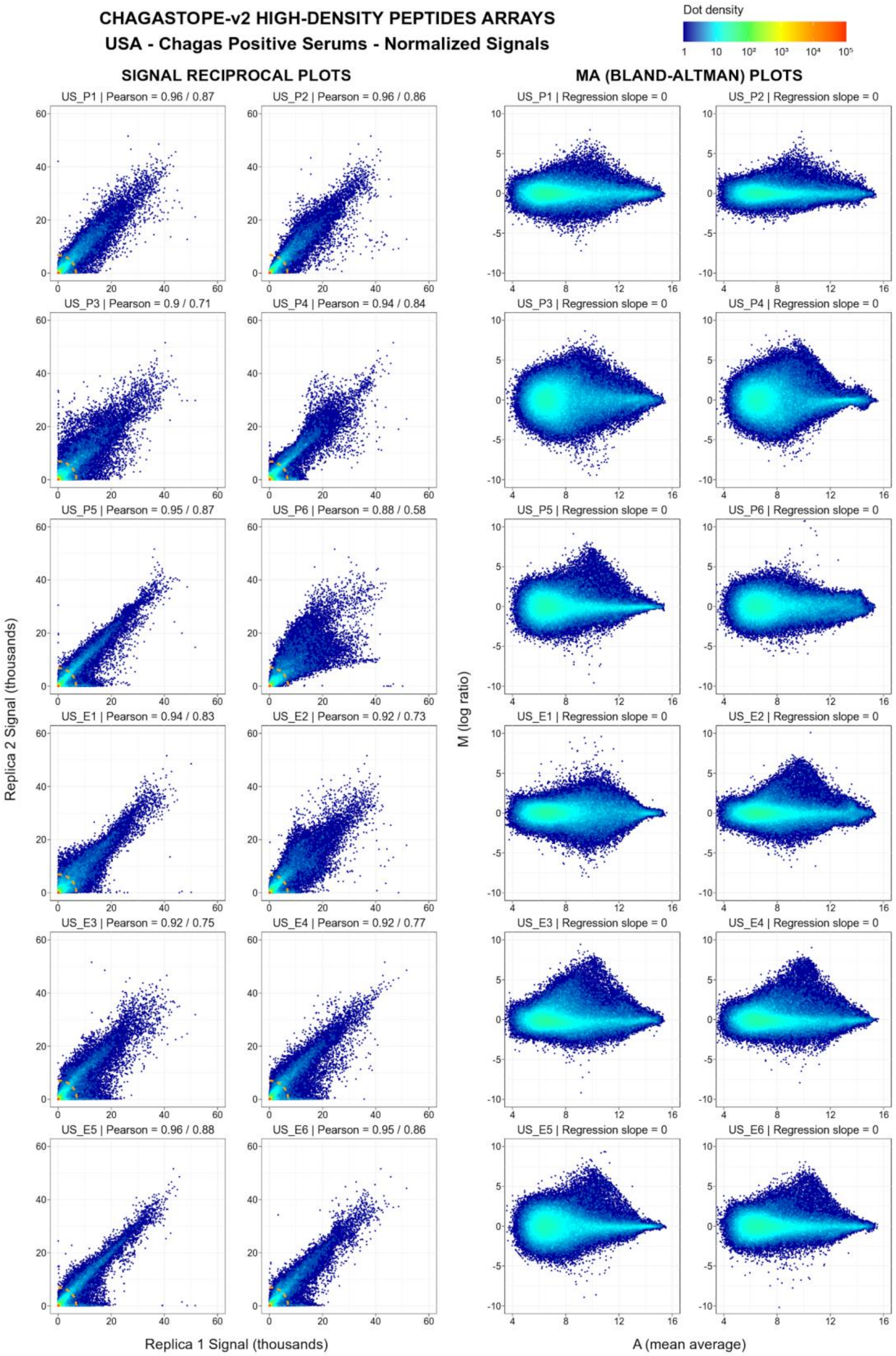


***Supplementary Figure S3l. Quality Control of 12-plex microarray assays for the US samples after normalization.*** *The legend is similar to that in Supplementary Figure S3a, but for the normalized signals.*

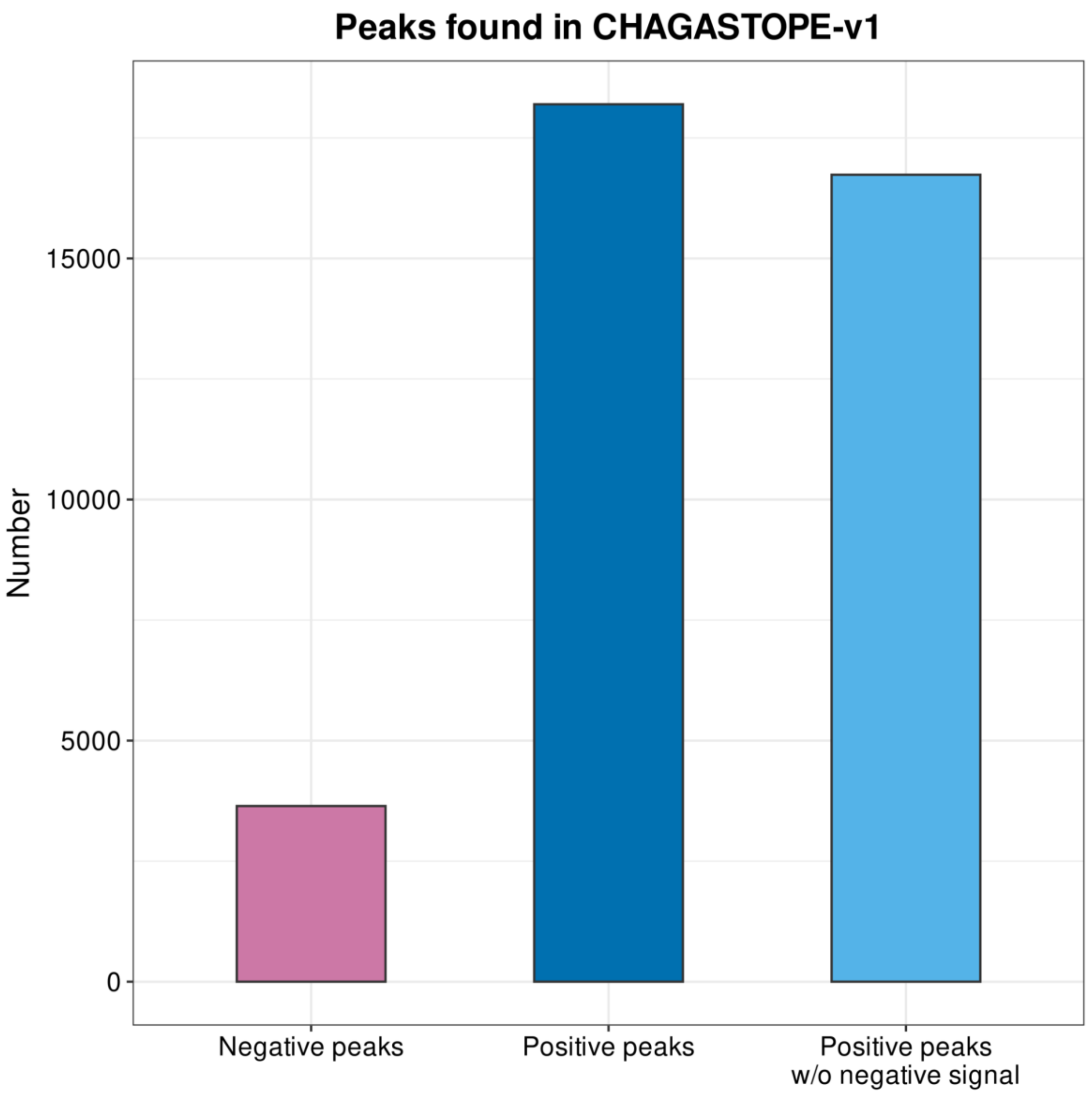


***Supplementary Figure S4. Number of antibody-binding peaks in proteins.*** *Comparative view of the reactivity of pools of Chagas-negative samples from healthy subjects and those from Chagas-positive subjects against T. cruzi proteins (summarized as peaks as per the definition in main text and Methods). The rightmost column shows the number of peaks in Chagas-positive subjects that show no significant signal in the negative samples (see Methods for details).*

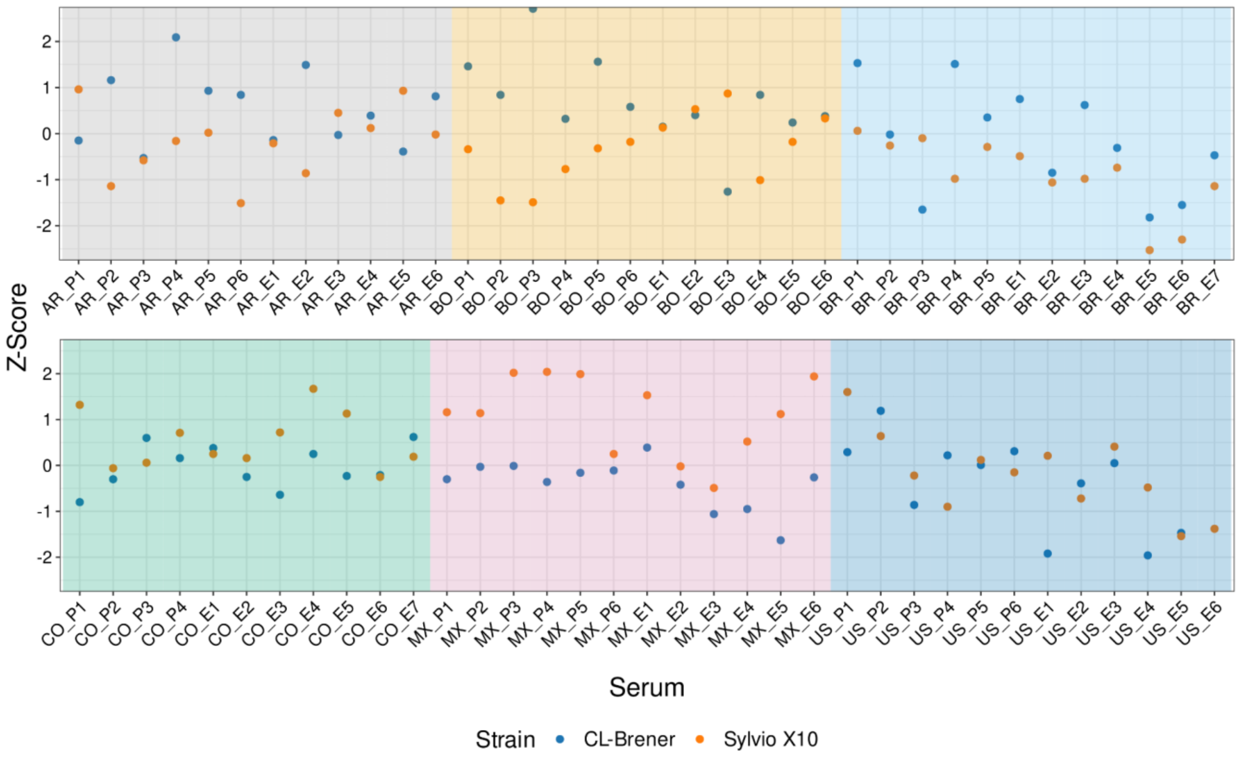


***Supplementary Figure 5. Comparison of strain-specific antigenicity across subjects.*** *Counts of reactive peptides per subject (those above the antigenicity threshold) in the two analyzed T. cruzi strains were standardized using z-scores. Standardization was necessary because CL-Brener and Sylvio X10 strains have different numbers of encoded proteins. Z-scores above 0 and below 0 represent higher and lower relative reactivity, respectively, across a given genome. This figure is similar to Figure 5 in the main paper but includes data from all analyzed subjects. Interpretation of these data must be done with care (caveat emptor) because some samples (those containing "_E" in the sample name) were not used for antigen discovery and selection, hence we may not be measuring limited peptide reactivity in these cases.*

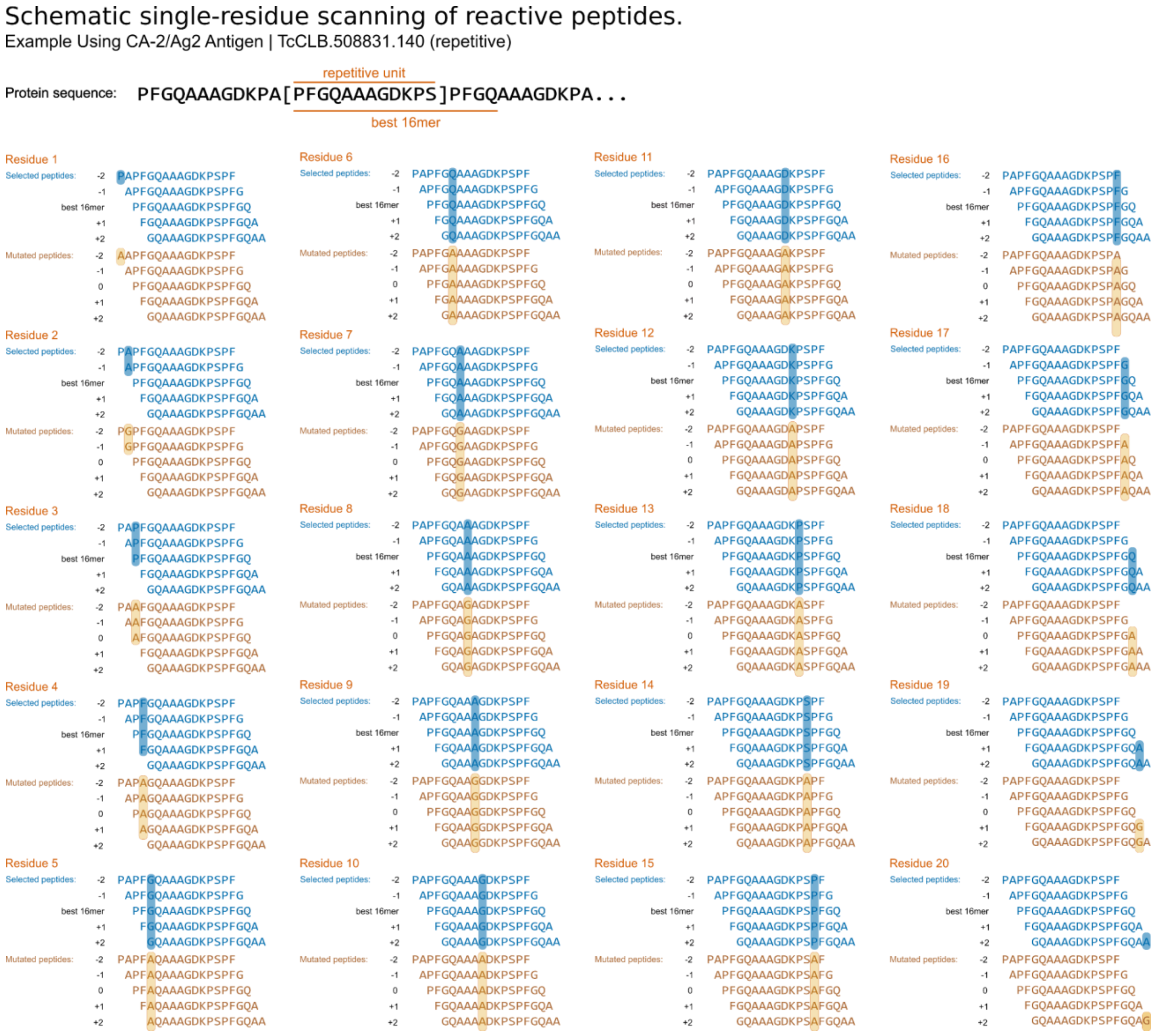


***Supplementary Figure S6. Complete visualization of single-residue mutagenesis scanning of epitopes.*** *Mutational scanning assessment scheme for one analyzed epitope. The same procedure was used to perform single-residue mutagenesis for other epitopes. Antibody-binding (reactivity) was measured for 1 to 5 original (non-mutated) peptide sequences (in blue), and for 1 to 5 mutated peptides per residue position (in orange). In all cases the central peptide (at position 0 in the figure) corresponds to the best 16mer (higher signal) in the epitope. Additional peptides positioned at -2, -1, +1, and +2 were used to measure the effect of mutations in different relative positions within a given epitope. Thus, for any given selected 16mer, a total of 80 mutated peptides were analyzed. Mutations were always to an Alanine, except where the original residue was an Alanine (it was substituted for a Glycine in these cases).*

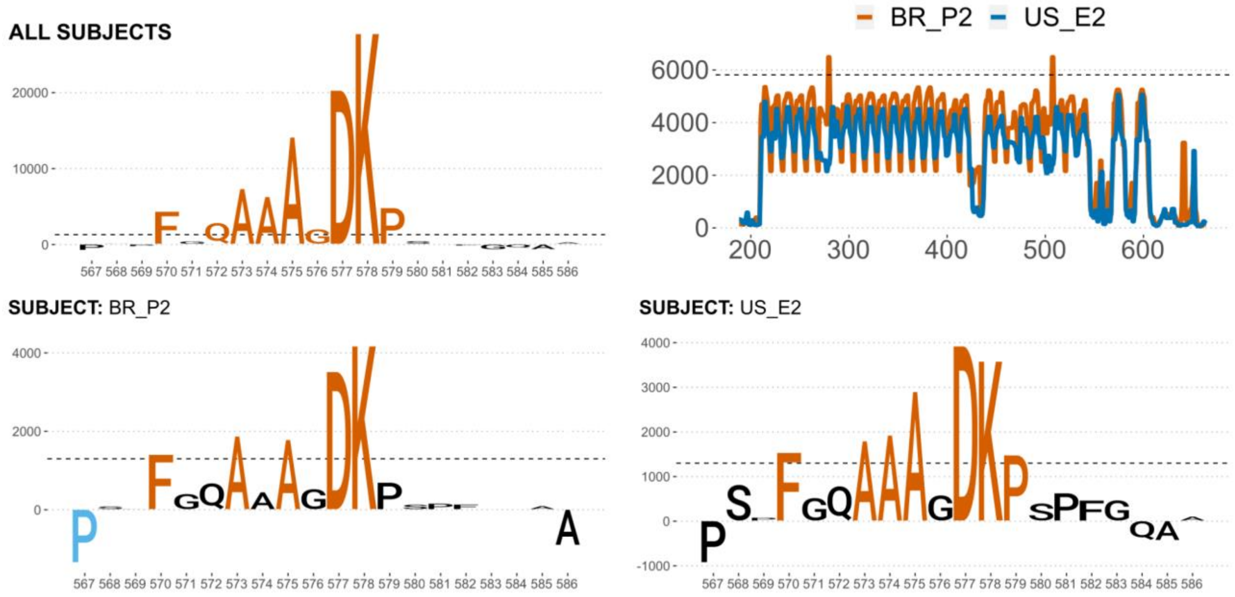


***Supplementary Figure S7. Single-residue mutational scanning of the Ag2/CA-2 antigen for subjects that were negative or displayed low signal against this epitope.*** *Top left, sequence logo summarizing key residues revealed by mutational scanning (derived from analysis of all subjects, logo is the same as in Fig 6 in the main manuscript). Top right, antibody-binding profile for the BR_P2 and US_E2 subjects and signal threshold (dashed line). Bottom, sequence logos of two subjects that display low signal and/or are negative with the chosen signal threshold.*

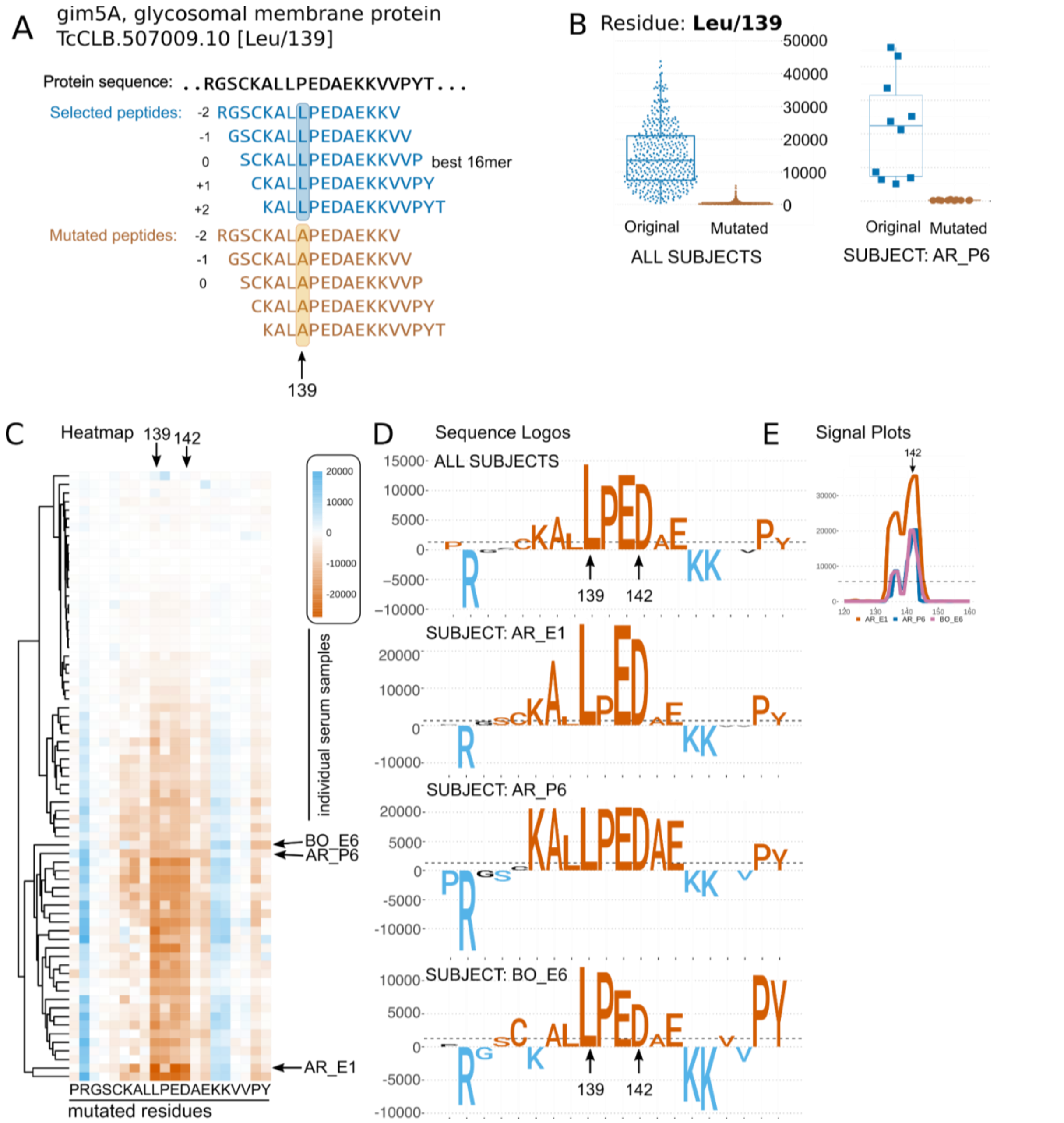


***Supplementary Figure S8. Single-residue mutational scanning of the novel Gim5A antigen.*** *The single antigenic peak in Gim5A was subjected to a mutational scanning as described (see main text).* ***A)*** *Schematic representation of the mutational scanning procedure, shown for one residue only (Leu139), for clarity.* ***B)*** *Average signal of the mutated vs wild-type peptides for all subjects and for one selected subject (AR_P6).* ***C)*** *Heatmap summarizing the mutational scanning for all residues and all subjects. The heatmap shows signal change from original to mutated peptides). Mutations that decrease antibody-binding are shown in different shades of orange, while those that increase binding are shown in shades of blue. Columns = residue positions, Rows = individual serum samples.* ***D)*** *Sequence logos summarize data for all positive sera (top), or for individual cases as denoted (y-axis: signal change in arbitrary units). Colors follow heatmap.* ***E)*** *Antibody binding signal plots for the selected subjects.*

Detailed characterization of antigenic responses as observed in different individuals and against different antigens, led to the observation that in some cases (this is mentioned in the main text), individuals display variable responses against the same antigen. Here, we document these observations with detailed figures for two examples in Supplementary Figures S9 and S10 below. These figures show differential recognition of the same antigen by individuals. Detailed epitope characterization using single residue mutagenesis showed that the core epitope motifs recognized by these individuals are different. In both figures we show the overall antigen-binding (peaks) in part A of the figure, and a zoomed in section of one/two peaks in section B of the figure. Finally in sections C and D we showed the characterization of the epitopes using single-residue mutagenesis (alanine-scanning).

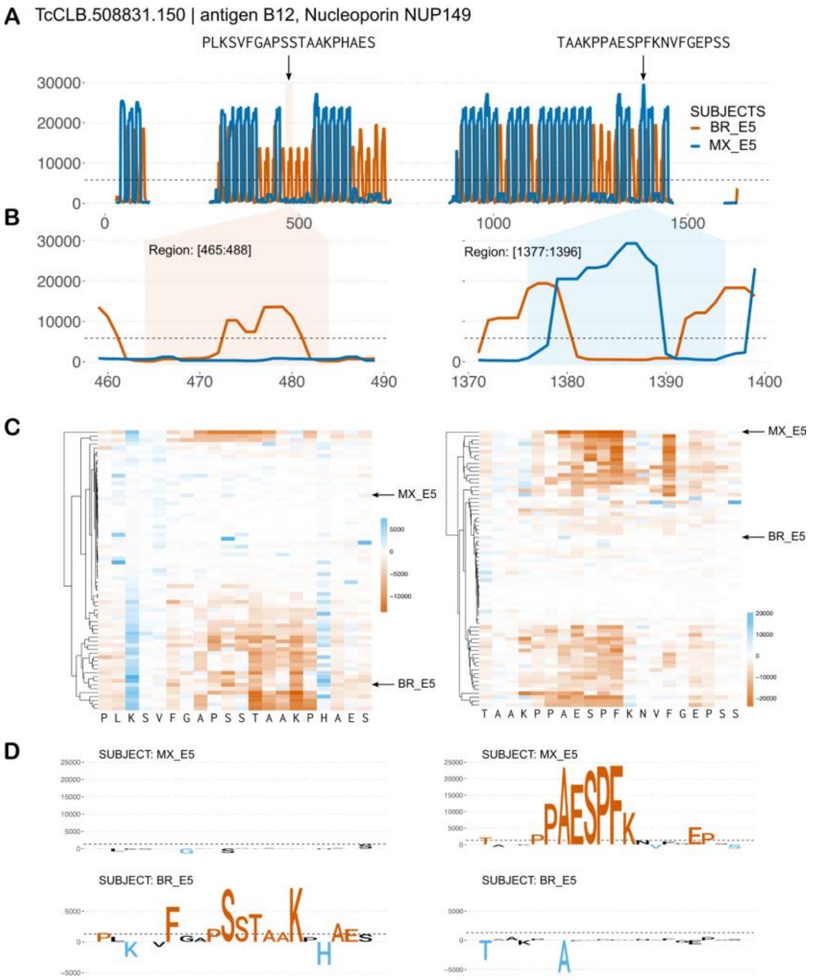


***Supplementary Figure S9. Single-residue mutational scanning for antigen B12 (TcCLB.508831.150).*** *Two sequences from the repetitive antigenic region of the B12 antigen (Gruber and Zingales, 1993) were subjected to mutational scanning as described (see main text).* ***A)*** *Individual antibody-binding signal profiles for two subjects; MX_E5 (blue) and BR_E5 (orange). The X axis represents the middle position of the peptide in the sequence of the protein. The Y axis represents the normalized signal (see methods for details) for each individual; regions without data were not present in the CHAGASTOPE-V2 design.* ***B)*** *Zoom in regions [458:490] (left) and [1370:1400] (right).* ***C)*** *Heatmaps summarizing the mutational scanning for all residues and all subjects (see legend of Fig 6).* ***D)*** *Sequence logos summarizing data for individual cases as denoted (y-axis: signal change in arbitrary units). Colors follow heatmap.*

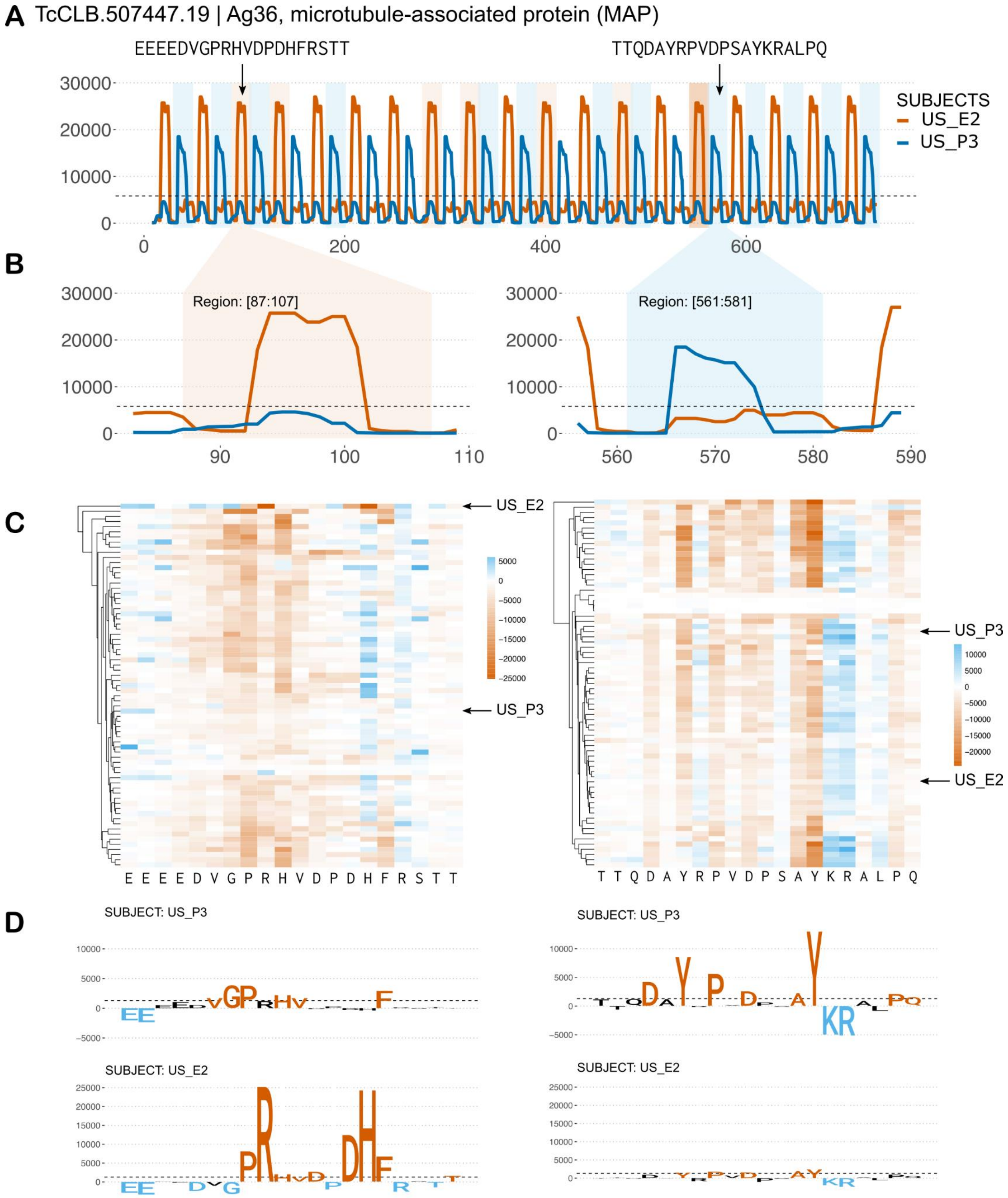


***Supplementary Figure S10. Single-residue mutational scanning for antigen Ag36 (TcCLB.507447.19).*** *Two sequences from the repetitive region of the Ag36 antigen (Ibañez et al., 1988, 1987) were subjected to mutational scanning as described (see main text). The legend is similar to that in Supplementary Figure S9. Subjects: US_P3 (blue) and US_E2 (orange) and zoom in regions [82:110] (left) and [555:590] (right).*

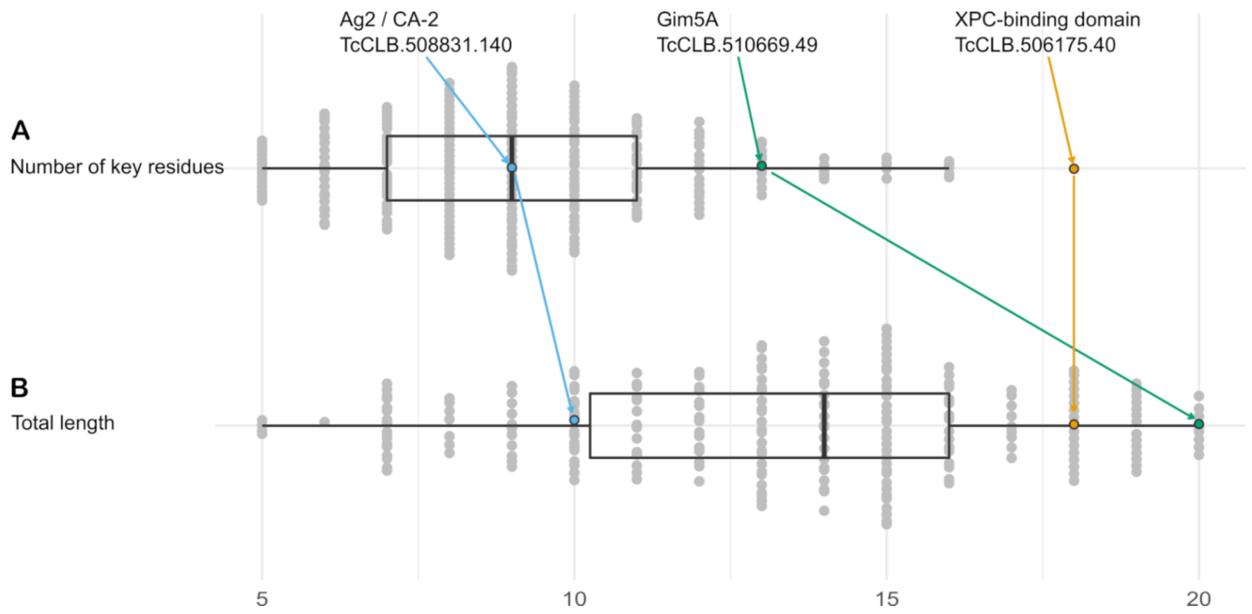


***Supplementary Figure S11. Characterization of core functional motifs from single-residue mutational scanning.*** *Summary of results for the 232 antigenic sequences analyzed with single-residue mutational scanning. A) Distribution of the number of key residues required for antibody binding (see Methods). The boxplot shows the distribution of values, average and most frequent number was 9. B) Distribution of total motif length, where motif length is the span from the first (leftmost) key residue in the motif to the last (rightmost). Average length span was 14, and most frequent length span was 15 residues. Three T. cruzi antigens are mentioned as examples.*

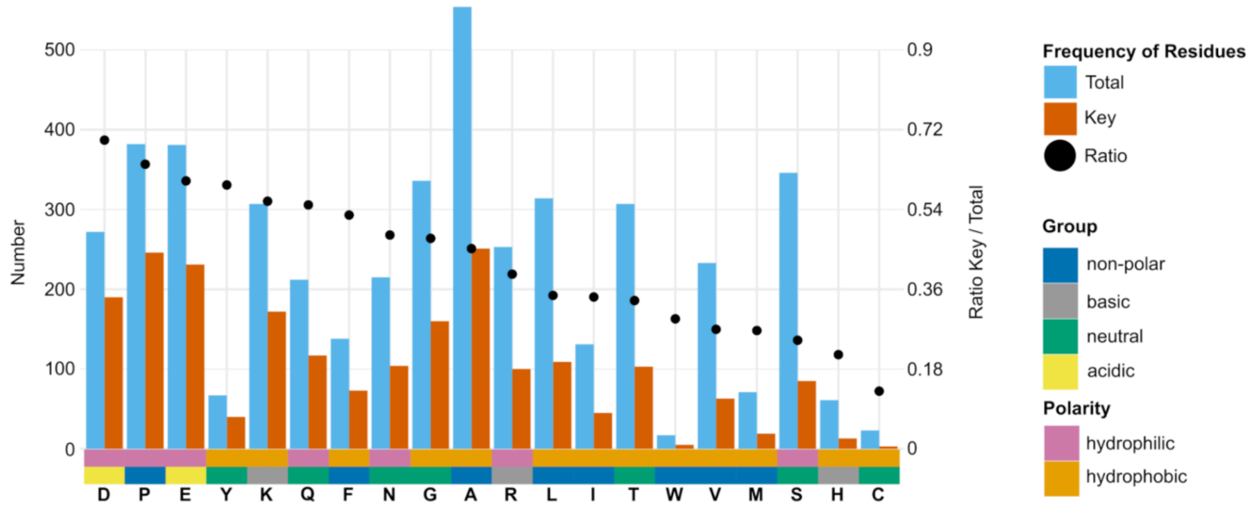


***Supplementary Figure S12. Characterization of core functional residues.*** *Distribution of amino acid residue types found in the 232 epitopes analyzed with single-residue mutational scanning (blue = all residues; orange = key residues important for antibody binding). Single black points represent the ratio of key / total residues (ratio = 1 if all residues are key residues). Amino acids are sorted based on this ratio from left (Asp/D, ratio ~ 0.7) to right. To aid the visualization, physicochemical properties of the amino acids are represented in colors, as shown.*

### Supplementary Information Guide

#### Tables

- **Tables available as spreadsheets: file *"Supplementary Tables.xlsx"***
    - Supplementary Table S1: Overview of arrays used in this work
    - Supplementary Table S2: Serum samples and pools
    - Supplementary Table S3: CHAGASTOPE array slides and assays
    - Supplementary Table S4: T. cruzi sequences cross-reactive with Leishmaniasis samples
    - Supplementary Table S5: T. cruzi sequences cross-reactive with normal human serum (healthy subjects, Chagas-negative)
    - Supplementary Table S6: Antigens reactive in all Chagas-positive pooled serum samples
    - Supplementary Table S7: Best antigenic region per cluster in individual serum samples
    - Supplementary Table S8: Single-residue mutational scanning results for shown epitopes
    - Supplementary Table S9: Detailed single-residue mutational scannings results for all epitopes
- **Tables available as standalone tab-delimited files (Figshare)**
    - **Supplementary Table S10. Mapping of CHAGASTOPE-v1 data to *T. cruzi's* proteins. Size ~ 62 MB.**
      TSV file containing the detailed information to map each peptide in CHAGASTOPE-v1 microarray to their original spots in each protein of *T. cruzi* in this array.
    - **Supplementary Table S11. Mapping of CHAGASTOPE-v2 data to *T. cruzi's* proteins.**
      TSV file containing the detailed information to map each peptide in CHAGASTOPE-2 microarray to their original spots in each protein of *T. cruzi* in this array.
    - **Supplementary Table S12. Detailed antigenic region analysis for pooled serums.**
      TSV file containing the detailed information for all the antigenic regions found in CHAGASTOPE-v1 and their respective antigenicity in the pooled serums.
    - **Supplementary Table S13. Detailed antigenic region analysis for individual serums.**

TSV file containing the detailed information for all the antigenic regions found in CHAGASTOPE-v1 and their respective antigenicity in the individual serums.

**Files (Figshare)**

- **Supplementary File S1. CHAGASTOPE-v1 positive controls.**
  PDF with the protein profiles for the positive controls analyzed in CHAGASTOPE-v1, showing their antigenicity in the pooled serums.
- **Supplementary File S2. CHAGASTOPE-v1 profiles for proteins above 3SD. Size ~ 2GB**
  ZIP file containing 3 PDF files with the protein profiles for the proteins in both *T.cruzi* proteomes analyzed in CHAGASTOPE-v1, showing their antigenicity in the pooled serums. Only proteins that had at least one peptide above an antigenicity threshold of mode + 3 standard deviations are shown.
- **Supplementary File S3. CHAGASTOPE-v2 profiles for proteins above 2.4SD. Size ~ 360 MB**
  ZIP file containing a PDF with the squashed protein profiles for the antigenic regions analyzed in CHAGASTOPE-v2, showing their antigenicity in the individual serums.
- **Supplementary File S4. CHAGASTOPE-v2 profiles for regions above 2.4SD. Size ~ 264 MB**
  ZIP file containing a PDF with the squashed region profiles for the antigenic regions analyzed in CHAGASTOPE-v2, showing their antigenicity in the individual serums.
- **Supplementary File S5. CHAGASTOPE-v1 & CHAGASTOPE-v2 smoothed signals. Size ~ 965 MB**
  2 ZIP files containing several TSV files (one per serum) with the parsed signal data (normalized and then smoothed) for each peptide in each protein in CHAGASTOPE-v1 and CHAGASTOPE-v2.
- **Supplementary File S6. Detailed Alanine Scan for Ag2 antigen.**
  PDF file containing the detailed scheme of single residue mutational scanning of CA-2/Ag2 Antigen (TcCLB.508831.140) alongside results for one serum as example.
- **Supplementary File S7. Alanine Scan general results.**
  PDF file with a summary of the single residue mutational scanning experiment results for each antigen. It shows a heatmap with residue relevance for all sera and 4 sequence logos (one for all positive sera and 3 other individual examples).
- **Supplementary File S8. Alanine Scan results for repeats.**
  PDF file with single-residue mutagenesis of repetitive *T. cruzi* antigens and the

multiple sequence alignment of repeats, as identified by Xstream (Newman and Cooper, 2007).
